# Sex differentiation, divergence, and introgression in the leishmaniasis vector *Lutzomyia longipalpis*

**DOI:** 10.1101/2025.04.08.647763

**Authors:** Adam MM Stuckert, David Turissini, Emmanuel RR D’Agostino, David Peede, Maya Parvathaneni, Jonathan A. Rader, Carlos Muskus, Eduar Bejarano, Jeremy R. Wang, Rafael Vivero, Daniel R. Matute

## Abstract

An estimated one billion people are at risk of acquiring the disease leishmaniasis annually. This disease, among others, is transmitted to humans via sandfly vectors of the family Psychodidae. In the Neotropics, *Lutzomyia longipalpis* is one of the most common vectors of leishmaniasis, but little is known about its genetics and population structure, hampering control efforts. We generated sex-specific genome assemblies of *Lu. longipalpis* from Colombia, and examined population genomics of this species. We found that sex accounts for ∼11% of genome-wide variation, primarily driven by differentiation between the sexes in an XY sex chromosome. Analyses of transposable elements indicate that the size difference in the sex chromosome is driven by more transposable elements in the male chromosome. Further, we examined population genomics and structure across the South American continent. We identified significant genetic divergence between Colombian and Brazilian sandflies and population structure within *Lu. longipalpis sensu lato* samples collected from the same sites in Brazil. These results strongly suggest the presence of multiple, sympatric species within a cryptic species complex, concordant with other published phenotypic evidence.

**SIGNIFICANCE:** Leishmaniasis has emerged as an important public health issue. Transmission occurs exclusively through Phlebotomine sandflies of the genera *Lutzomyia* and *Phlebotomus*, yet our ability to prevent its spread is constrained by limited knowledge of the biological and ecological factors that make these insects effective vectors. In this manuscript, we leverage whole-genome sequencing to investigate the evolutionary forces shaping genetic diversity in *Lutzomyia longipalpis*, a key vector of leishmaniasis.

## INTRODUCTION

Phlebotomine sandflies of the genera *Lutzomyia* and *Phlebotomus* are the only known vectors of *Leishmania*, a protozoan parasite that is transmitted to humans^1,2^. *Leishmania* infections can lead to the clinical disease leishmaniasis, which can cause skin ulcers and scarring, destruction of mucous membranes, extreme organ damage (e.g., liver, spleen, and bone marrow), and can be fatal in some cases. *Leishmania* is endemic to 98 countries and an estimated one billion people are at risk of infection^3,4^. Despite its high prevalence and disease burden, leishmaniasis is poorly understood, and thus is classified as a neglected tropical disease^5,6^. Although leishmaniasis is largely regarded as a tropical disease, *Lutzomyia* ranges are expanding northward^7^ and the U.S. is experiencing increasing caseloads of leishmaniasis^7–9^. Ecological niche modeling predicts that leishmaniasis is likely to become more common in the U.S. due to climate change^10^, and this is already evident via increasing clinical cases^11^. Treating leishmaniasis is difficult and often prohibitively expensive for individuals in underserved areas with endemic *Leishmania,* thus, preventing the spread of leishmaniasis is critically important^12^. Perhaps the most effective way to prevent the transmission of vector-transmitted diseases like leishmaniasis is to control the vector itself^13^. Despite the clear importance of *Leishmania’*s vectors for human well-being, preventive efforts are hampered by our lack of knowledge of the genetic and ecological factors that make sandflies effective vectors of *Leishmania*. Compounding the problem, functional studies of leishmaniasis vector biology are hindered by a variety of gaping holes in scientific knowledge, including: taxonomy and phylogenetics of sandflies, genetic characteristics of sandfly susceptibility to leishmaniasis, and a lack of genomic resources.

In the Neotropics, there are ∼40 *Lutzomyia* species that transmit *Leishmania*, but one of the most commonly implicated and widespread is *Lutzomyia longipalpis.* This species has a range from Argentina through Mexico^2,14^. Complicating our understanding of this species’ biology, a variety of lines of evidence suggest that *Lu. longipalpis* is actually a species complex containing a number of cryptic species with substantial phenotypic divergence across its geographic range^15,16^. This evidence includes variation in microsatellites of key genes^17–19^, courtship song^20^, morphology^21,22^, and pheromones^18^. Nevertheless, this “species” is thought to be the major vector of *Leishmania infantum,* the causative agent of leishmaniasis in the Neotropics^23^. This species inhabits a wide variety of microhabitats, including silvan habitats and inside of rural houses^24–29^. Like most phlebotomine sandflies, females of this species are hematophagous (feed on blood), and can spread leishmaniasis to birds, dogs, pigs, horses, humans, and a variety of other mammalian hosts during feeding, meaning that common subsistence animals or pets can be intermediate hosts and promote vector populations^30^. Dramatic environmental changes in human land use (e.g. deforestation, roads, etc.) have caused concomitant changes in *Lu. longipalpis* distribution and habitat use. Of particular note is that *Lu. longipalpis* has shown a shift in recent decades to live and breed in highly populated urban areas^27^. Given that this species is a common vector of leishmaniasis, this puts an ever-increasing number of people at risk of acquiring leishmaniasis.

In this piece, we address fundamental genetic questions in the widespread leishmaniasis vector *Lutzomyia longipalpis.* First, we present highly-contiguous chromosomal and sex-specific genome assemblies of *Lu. longipalpis.* We then use these new assemblies as tools to examine population genomics of this sandfly from the Andean region of Colombia. We then further use publicly available genomic data from Brazilian *Lu. longipalpis* to examine population genomics of this vector species across the South American continent. As predicted by previous work suggesting *Lu. longipalpis sensu lato* is a cryptic species complex, we find extensive population differentiation between Colombian and Brazilian populations. Additionally, our population genomic analyses indicate extremely low nucleotide diversity, elevated runs of homozygosity, and lower effective population sizes in Colombia relative to any Brazilian population. Our results provide a roadmap for the implementation of genomic analyses in sandflies and to ultimately understand the molecular and genetic underpinnings of the interactions between vectors and parasites.

## RESULTS

### Genome sequencing

We produced sex-specific genomes for female and male Colombian sandflies (Table 1). Table S1 lists all the accession numbers. The initial female contig assembly built from PacBio HiFi reads was 148 Mbp in length spread across 2,254 contigs and had an N50 of 1.25 Mbp. The male contig assembly built from PacBio HiFi reads was 159.7 Mbp in length spread across 1,835 contigs with an N50 of 1.14 Mbp. The inclusion of HiC data improved our genome assembly. Scaffolding our assemblies with HiC data yielded putative chromosome-level assemblies for both female and male sandflies. The female assembly was 151.2 Mbp in length and contained four chromosomes with lengths of 47.7, 37.2, 29.0, and 22.8 Mbp and represented over 95% of the total genome. The female assembly contained >95% of complete dipteran orthologs: 94.16% in complete single copies, 1.13% in complete duplicated copies, 0.46% fragmented, and 4.26% missing. The male assembly was 159.7 Mbp in length and contained four chromosomes of 49.0, 36.8, 29.3, and 22.5 Mb, representing over 95% of the total genome. The male assembly contained just under 95% of complete dipteran orthologs: 93.49% in complete single copies, 1.19% in complete duplicated copies, 0.33% fragmented, and 4.99% missing.

**Table 1.** Genome statistics. The Brazilian assembly comes from^31^, and is derived from a colony collected in the 1980s from Jacobina, Bahia State, Brazil. Our female and male assemblies are derived from wild-caught sandflies from Ricaurte, Colombia.

|  | Brazilian *Lu. longipalpis* assembly | Female contig | Female scaffolded | Male contig | Male scaffolded |
| --- | --- | --- | --- | --- | --- |
| Summary |  |  |  |  |  |
| Assembled bases (Mbp) | 159.7 | 1997 | 151.2 | 159.7 | 159.7 |
| Number of contigs | 383 | 151.2 | 2002 | 1835 | 1849 |
| Contig N50 (Mbp) | 1.04 | 1 | 1 | 1.1 | 1.1 |
| Number of scaffolds | 138 | - | 1742 | - | 1618 |
| Scaffold N50 (Mbp) | 40.6 | - | 37.2 | - | 36.8 |
| Percentage of assembly in scaffolds | 93.2 | - | 91 | - | 86.7 |
| BUSCO (percentages) |  |  |  |  |  |
| Single copies | 90.5 | 94.12 | 94.16 | 93.9 | 93.49 |
| Duplicated copies | 6.73 | 1.13 | 1.13 | 1.19 | 1.19 |
| Fragmented copies | 0.33 | 0.46 | 0.46 | 0.4 | 0.33 |
| Incomplete copies | 0 | 0 | 0 | 0 | 0 |
| Missing | 2.89 | 4.29 | 4.26 | 5.02 | 4.99 |
| RepeatMasker (Mbp) |  |  |  |  |  |
| Percentage of repeats (%) | - | - | 15.69 | - | 15.4 |
| Repeat bases (Mbp) | - | - | 23.7 | - | 24.6 |
| Retroelements | - | - | 4.37 | - | 4.92 |
| DNA transposons | - | - | 19.3 | - | 19.6 |
| Small RNA | - | - | 0.39 | - | 0.0319 |
| Simple repeat | - | - | 1.3 | - | 1.29 |
| Low complexity | - | - | 0.47 | - | 0.5 |
| Total bases masked | - | - | 25.69 | - | 26.67 |

Next, we identified the differences between the male and female genomes (Figure 1A). We identified 640,461 unique kmers associated with sex-limited chromosomes using SRY on our Illumina data. These kmers mapped to 111,862 HiFi reads spanning 888 Mbp in the male dataset. Our initial draft “Y” chromosome assembly was 26.7 Mbp across 740 contigs. After scaffolding, we placed 691 contigs (93.4% of contigs) into 50 scaffolds in an assembly that was 26.8 Mbp in length (including placed gaps) with a largest scaffold of 26.2 Mbp (thus the other 49 scaffolds contain ∼0.6 Mbp). Thus, we were able to scaffold nearly the entire section of the “Y” chromosome that is limited to males into a single scaffold, with minimal unplaced content outside of this scaffold.

**Figure 1.**
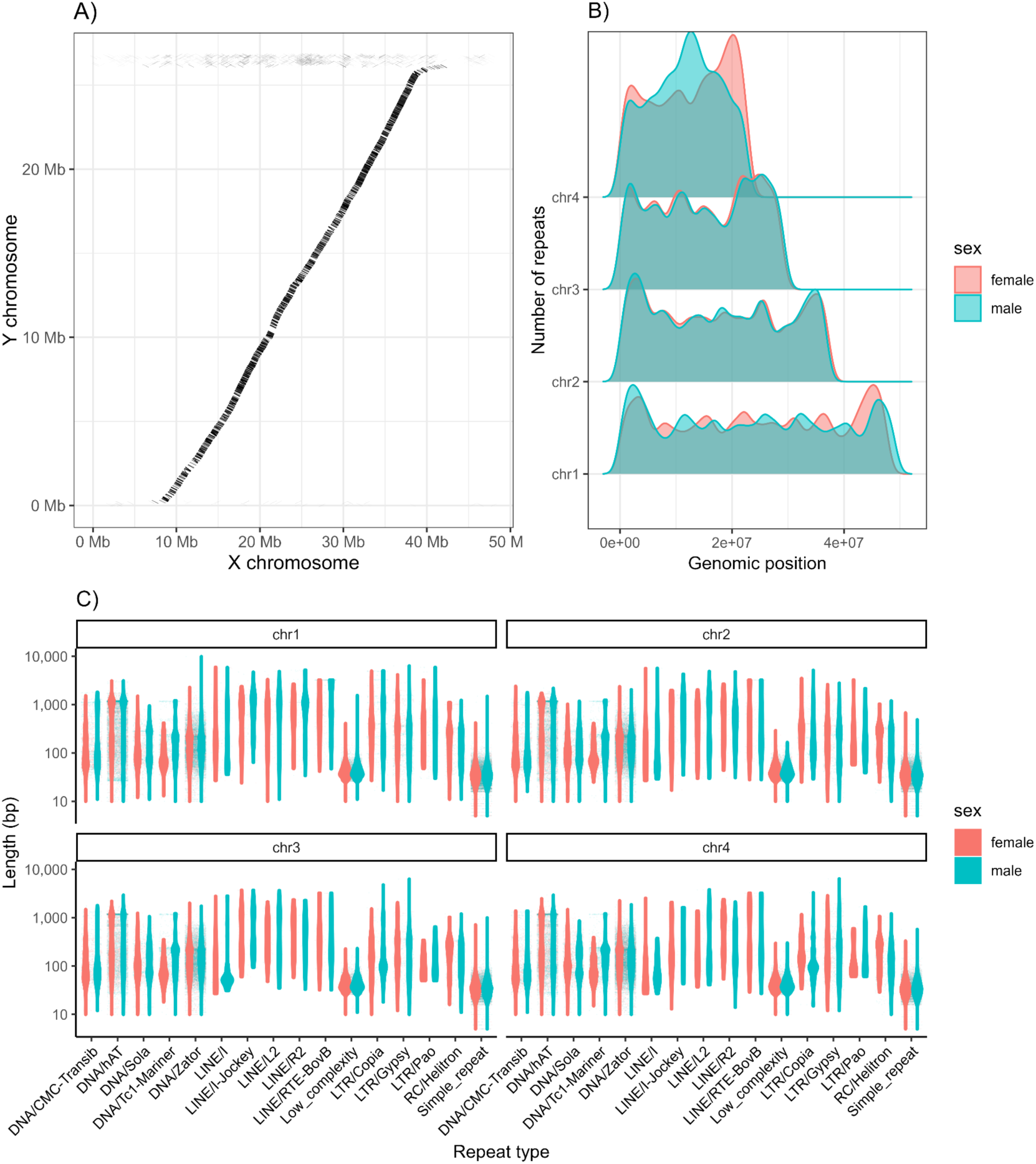
The sex chromosomes of *Lu. longipalpis*. **A.** Synteny plot of alignments between assembled male (presumptive) X and Y chromosome sequences. **B.** Number of repeats in the female and male genome assemblies based on genomic position. **C.** Violin plots of TEs in the female and male genome assemblies and their length.

### Comparison between the sexes indicates increased transposable element content in males on the sex chromosome

Transposable elements (TEs) are considered important drivers of sex chromosome evolution^32,33^, so we quantified transposable element content in the female and male assemblies. RepeatMasker identified and masked a total of 16.99% (female) and 16.70% (male) of the genome, 15.69% and 15.4% of which were TEs in the female and male assemblies respectively (Figure 1B). Most repeats were DNA transposons (12.8% and 12.3% of the genome in females and males respectively) or Retroelements (2.9% and 3.1% of the genome in females and males respectively, Figure 1C). A small proportion of the genome were simple repeats (0.9% and 0.8% of the genome in females and males respectively). The difference in relative proportion of transposable elements between the sexes is largely driven by chromosome 1. This chromosome is not only longer in males (49 Mbp vs 47.7 Mbp), but also has a larger overall proportion of bases masked in males (15.89% vs 13.60%, Figure 1C). The male chromosome 1 has a total of 7.8 Mbp of transposable elements masked by our repeat masking pipeline, whereas the female chromosome has 6.499 Mbp of transposable elements masked. The difference, 1.3 Mbp, corresponds to the total difference in size between the assembled male and female chromosome 1 (see Figure S1). In the three autosomal chromosomes, the proportion of transposable elements to assembled chromosome length is <0.5% different between females and males.

### Genome annotation

We annotated both sexes of our Colombian *Lu. longipalpis.* In total, we annotated 14,559 genes in the male assembly; 10,874 of these predicted genes were in the four chromosomes (4008, 2607, 2371, and 1888 in chromosomes 1-4 respectively), as opposed to in the contigs. In the female assembly, we annotated 12,029 genes in total; 10,934 of these predicted genes were in the four chromosomes (3,922; 2,662; 2,371; and 1,979 in chromosomes 1-4 respectively). The predicted genes are approximately equal to RefSeq predictions in *Lu. longipalpis* from other localities (GCF_024334085.1), which has 12,045 predicted genes. In both sexes of our assemblies, the majority of predicted genes are found in the chromosomes. However, our male assembly has approximately 2,000 more predicted genes than existent assemblies. This is due to the presence of multiple copies of genes between the heteromorphic chromosomes — most of which are represented as unplaced contigs. While females had more predicted genes overall in the regions of the genome that are located in the chromosomes, males had a greater number of predicted genes owing to the predicted genes found in un-scaffolded contigs. Further, males had more predicted genes in chromosome 1, the heteromorphic chromosome. We identified a total of 3,462 genes annotated in the highly differentiated region of chromosome 1 (average F_ST_ > 0.3 between females and males) in the male assembly. Of these, we tentatively assigned identifications to 3,611 genes based on protein evidence from OrthDB version 11 and VectorBase version 67. Among the genes found in this region of high differentiation between females and males within our Colombian population are a number of genes related to courtship (e.g., *takeout*), sex-differentiation (e.g., *sex-lethal, fruitless, doublesex-and mab-3-related transcription factor A2, lethal(2)*), testes-specific genes (e.g., ejaculatory bulb-specific protein 3, ejaculatory bulb-specific protein 3-like, testis-expressed protein 10 homolog, motile sperm domain-containing protein), putatively related to feeding and finding mates (e.g., gustatory receptor genes, gustatory receptor for sugar taste 64f, odorant receptor genes), or related to blood meals (e.g., venom allergen 5).

Nucleotide diversity across the genome is similar in females (*π* = 0.00177 ± 0.00233 SD female assembly; *π* = 0.00170 ± 0.00228 SD male assembly) and males from Colombia (*π* = 0.00171 ± 0.00236 SD female assembly; *π* = 0.00159 ± 0.00225 SD male assembly; Figures 2A-2B). However, concordant with other results, nucleotide diversity was lower on chromosome 1 in female sandflies (*π* = 0.00138 ± 0.00192 SD female assembly; *π* = 0.00131 ± 0.00196 SD male assembly) than males (*π* = 0.00165 ± 0.00239 SD female assembly; *π* = 0.00160 ± 0.00225 SD male assembly).

**Figure 2.**
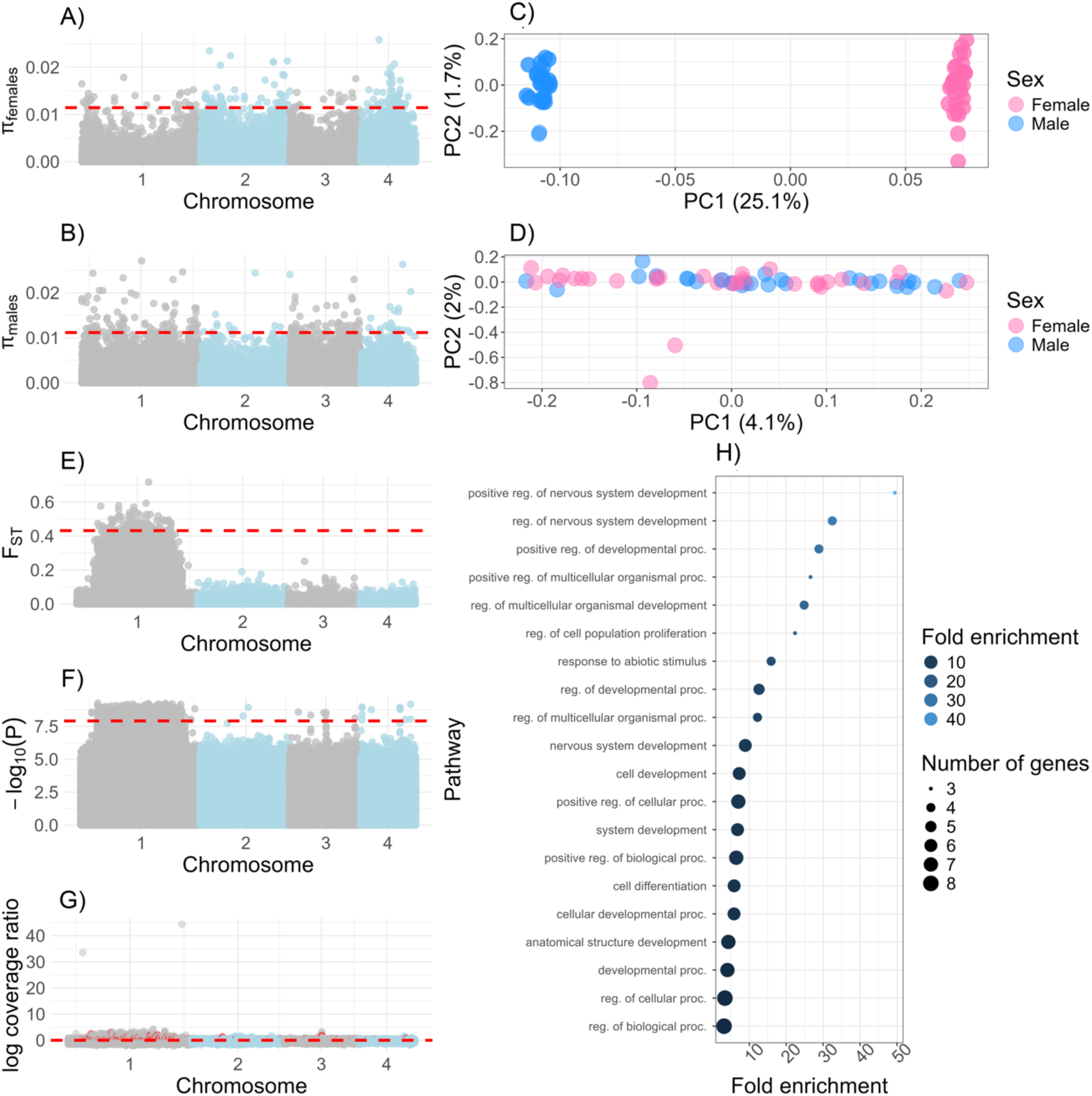
Sex chromosome differentiation in *Lu. longipalpis*. **A.** *π* in female sandflies from Colombia. **B.** *π* in male sandflies from Colombia. **C.** PCA of only chromosome 1. **D.** PCA of the autosomes, chromosomes 2-4. **E.** F_ST_ between the sexes. **F.** Manhattan plot of genome-wide association study between the sexes in Colombian sandflies. **G.** Coverage represented as a ratio of average male depth divided by average female depth, calculated as a 2,000bp mean. Two regions in chromosome 1 have been removed that have average log coverage of >30 (with mean positions at 5847001 and 45239001). **H.** GO biological processes analysis from GWAS results.

Next, we studied the extent of differentiation between the *X*-chromosome and the potentially male-restricted chromosome using PCA. We saw evidence of genomic differentiation between female and male genomes in our study. PC1, which corresponds to the sex of the sample, explained ∼11.2% of genome-wide differentiation. This signal was clearly driven by chromosome 1, which showed a similar pattern in which sex explains ∼25.1% of variance in the PCA (Figure 2C). This signal was entirely absent when we did PCA for the putative “autosomes”, chromosomes 2-4, with comparatively very little of the variation explained (PC1: 4%, PC2: 2% Figure 2D) and no correspondence to the sex of the sample. Genealogies based on chromosome 1 showed a similar result (Figure S2). Additionally, we saw a long, extended region of differentiation (i.e., elevated F_ST_) that occurred between female and male sandflies on chromosome 1 (Figure 2E). This region also shows an extended region of decreased nucleotide diversity in female sandflies (Figure 2A), but there was no concomitant decrease in nucleotide diversity within that region in male sandflies (Figure 2B). Given this finding, we compared *π* between sexes, with the specific hypothesis that *π* was lower on chromosome 1 in females than on autosomes. A linear model indicated that sex (*t* = 0.744, *p* = 0.4821) and the interaction of sex*chromosome (*t* = -0.984, *p* = 0.4821) were not a significant explanatory variable, but that chromosome was (*t* = 4.260, *p* = 0.381), and it was driven by lower *π* in the sex chromosome.

### Genome wide association studies revealed autosomal genes associated with sex

Next, we did a GWAS to identify genetic regions associated with sex determination. We identified 2,641 genic SNPs, and 3,165 intergenic SNPs associated with sex, the vast majority of which were located in the sex chromosome. The highly differentiated region between sexes in chromosome 1, the same that shows differentiation by our other analyses, showed a strong association with sex (Figure 2F). Nonetheless, there were autosomal markers strongly associated with sex. Sixty-seven SNPs were also associated with significant differences in genomic coverage between the sexes. Eight genic, autosomal SNPs mapped to three predicted genes in the three autosomal chromosomes, one of which has no detectable ortholog in any other dipteran (g5391). A second gene (g7272) showed strong similarity to *mdg3* retrotransposons in *D. melanogaster* (FBgn0043882). This gene also showed a difference in genomic coverage between the sexes in our short read sequencing dataset, with a log2(male:female) coverage ∼0.78. The rest of markers showed a uniform distribution of coverage around 1 (Figure 2G). The third and final gene with genic SNPs associated with sex was Apoptotic protease-activating factor 1 (*apaf-1*), which plays a role in sex-specific cell apoptosis and male infertility^34,35^. Twenty-four intergenic SNPs were in the autosomes, and they are in close proximity to seven genes and three transposable elements. Table S2 lists their name, location, association *p*-value, and known orthologs in other dipterans. No gene involved in primary sex determination showed a significant association. Of note, we saw a significant enrichment for genes involved in regulation of nervous system development (Figure 2H, GO:0051962: 49.4X; GO:0051960: 32.5X, FDR ≤ 9.6 × 10^−4^). Table S3 lists other GO categories with their associated enrichment values.

### Population genomics across the continent

*Lutzomyia longipalpis* has been proposed to be differentiated into distinct populations and even separate species across South and Central America^15,36,37^. We used a two-pronged approach to study the population structure between our Colombian samples and samples of females from several Brazilian populations^31^. Notably, Sobral was split *a priori* by Labbé et al.^31^ into Sobral 1S and Sobral 2S. First, we used phylogenetic trees. An unrooted genealogy based on 100kb windows revealed a clustering determined by geographical location consistent with previous studies^31^. Jacobina and Sobral 1S were each formed by two distinct and unrelated clusters (Figure 3A-3B). We further examined gene concordance between populations as an estimate of gene flow. We created a single “population” tree (akin to a species tree) from our multiple individual sampling of both the whole genome, and of BUSCO genes. When forced to recapitulate the populations, we found poor gene concordance support for several branches, indicating poor alignment between individual gene trees and our consensus population tree (Figure S3). This phylogenetic discordance may occur in instances of recent divergence, incomplete lineage sorting, hybridization, and poor tree inference which may be occurring in this population tree^38^. This system may be an amalgamation of all of these factors. In summary, the two analyses were consistent and suggested significant differentiation between lineages from different locales, particularly between Colombian and Brazilian samples. Further, we observed evidence for differentiation between samples from the same locale in Jacobina, Sobral 1S, and Sobral 2S (Figure 3A) which indicated the existence of differentiated and syntopic lineages. Second, we used PCA to represent the genetic variation across the continent (Figure 3B). The first three principal components (PCs) explain 61% of the genetic variance of South American *Lu. longipalpis* (Figure S4). PC1 accounted for ∼34% of genome-wide variation, and appears to reflect differentiation between Colombia and Brazil, and between some Brazilian lineages (i.e., Lapinha cave; Figure 3C). PC2 and PC3 also capture considerable variance (16.4 and 10.6% respectively; see Figure S5). PC2 separated Marajo and Sobral 2S from Ricaurte, Sobral 1S, Laphina with Jacobina samples split between the two groups. Sobral 2S, Marajo, and Ricaurte are similar in PC3, but Jacobina and Sobral 1S are divided into two separate clusters from each locality.

**Figure 3.**
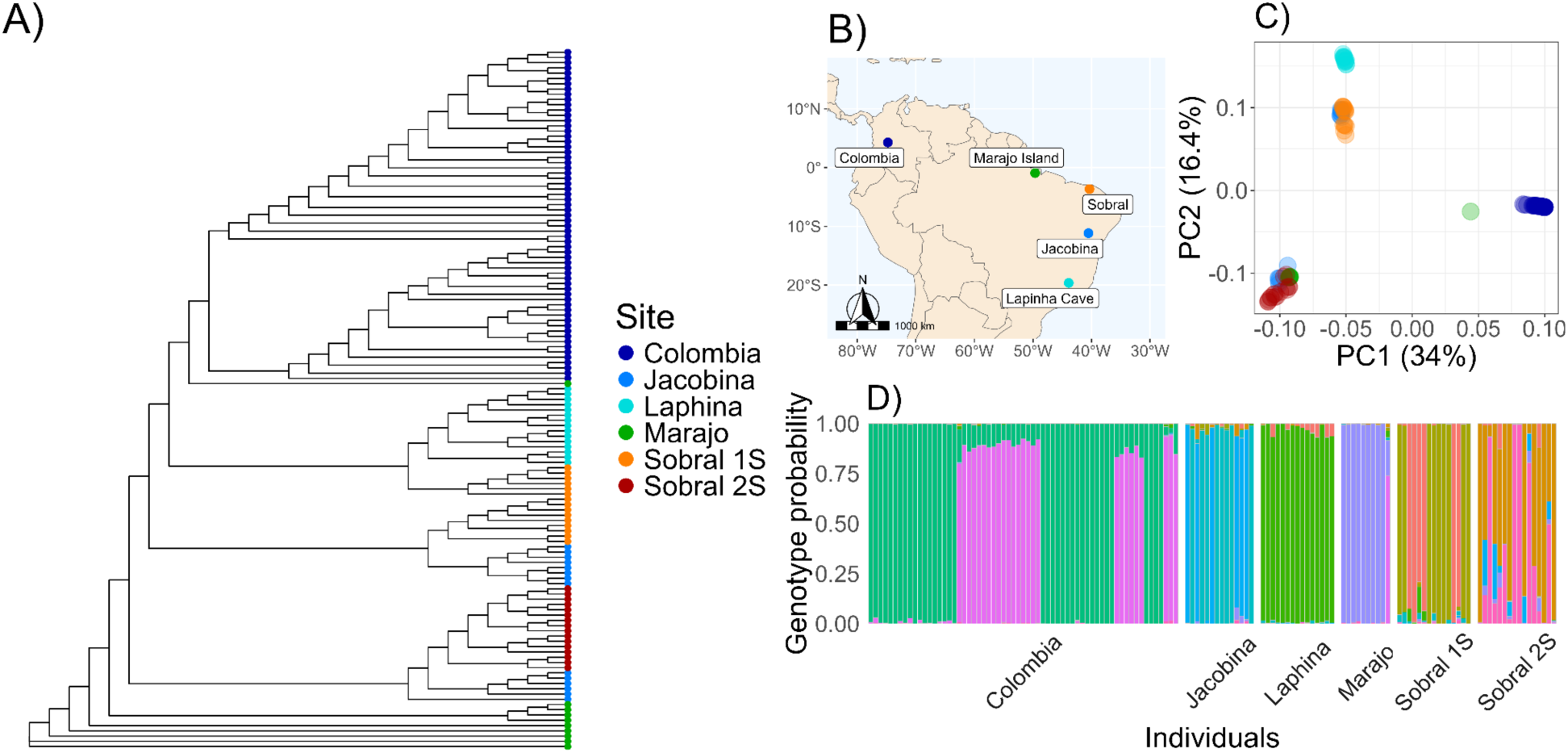
Partition of genetic diversity in Lu. longipalpis across South America. **A.** Phylogeny of sampled *Lutzomyia longipalpis* across South America. **B.** Sampling localities with genomic data included in population genetics analyses. **C.** Principal component analysis of Colombian sandflies from Colombian (this study) and Brazilian sandflies. **D.** Structure plot of genotype probabilities across these same samples. Colombian flies divide into two genotypes driven by sex. All Brazilian sandflies are females, while our Colombian samples also contain males. Panels A and C have the same legend for sampling locality, D has a different color schema due to the much larger number of genotypes predicted. Brazilian populations are from Labbe et al.^31^.

Next, we measured the extent of gene flow between these genetic clusters. Given the strong evidence of reproductive isolation between lineages^21,37^, we hypothesized reduced—or even nonexistent—gene flow between the populations we identified. We used *PCAngsd*, which identifies the most likely number of clusters, but also estimates admixture proportions from allele frequencies. Genetic variance was best explained by the existence of 10 clusters. Two of these clusters correspond to male and female *Lu. longipalpis* in Colombia, with little contribution to other lineages (a probability of <1% all Brazilian samples, except for the Colombian genotype that corresponds to males, which had a 73.9% probability in one Marajo sample, leading us to believe it was a male sandfly). This analysis (Figure 3D) also separated Colombian and Brazilian sandflies. The genetic variation in Brazilian samples was best explained by the existence of eight groups which correspond largely to the five geographic sites from which they were collected. The only exceptions were Sobral 1S, Sobral 2S, and Jacobina where we detected at least two main clusters per locality. The majority of individuals show limited evidence of admixed ancestry. In Colombia, Laphina, Jacobina, and all but one Marajo sandfly, admixture proportions are <10% per individual. This is particularly notable in the sites with syntopic lineages which could potentially have the chance to interbreed and yet do not appear to (Jacobina, some individuals within Sobral), as evidenced by divergent genotype clusters driving nearly the entirety of individual genetic makeup (e.g., blue vs teal individuals in Jacobina; Figure 3D). However, Sobral showed significant admixture within certain individuals (those of the “2S” lineage, Figure S6). We then proceeded to examine gene flow between populations using fbranch statistics. These analyses indicated high gene flow from Sobral 2S into Jacobina, Jacobina into Sobral 1S and Sobral 1S into Laphina (with Sobral “1S” and “2S” being *a priori* identified as different *Lu. longipalpis* lineages by^31^; Figure S7). Thus, there is either significant gene flow between populations or our consensus population tree does not adequately reflect cryptic diversification and potential syntopic lineages. Indeed, the consensus phylogeny of all individuals indicates the latter, as Sobral 2S individuals cluster with some Jacobina individuals. Overall, these results confirm that the different lineages of *Lu. longipalpis* are divergent, indicates advanced divergence towards speciation, and likely the presence of syntopic and hybridizing lineages.

The heterozygosity in the whole *L. longipalpis* clade was 0.06 (Figure 4A), but given the population structure described above, we studied the partitioning of genetic diversity within each *Lu. longipalpis* lineage. First, we quantified the extent of heterozygosity in each lineage (Figure 4B-E). Some of the populations are less genetically diverse than others. Colombia shows the lowest heterozygosity (0.0036); Sobral-1S and Marajo showed the highest heterozygosity (∼0.01) and were three times more diverse than Colombia. In general, the trends of heterozygosity along the genome were similar across populations (Figures 2A, 2B, 4F, 4G, Figure S8). All pairwise comparisons were significant (Figure 4H), indicating differences in the level of genetic diversity across *Lu. longipalpis* lineages. Second, we looked for runs of homozygosity (ROH), which are identical haplotypes inherited from each parent and thus are a measure of inbreeding^39^. The total length of ROH was correlated with the number of tracks in an individual (Figure 5A). Colombian *Lu. longipalpis* had more runs of homozygosity (ROHs) than any of the Brazilian populations (Figure 5B, C, Figure S9). The pattern is not driven by a single individual and is a property of the whole population (Figure S10) and extends beyond the mean level of linkage disequilibrium (Figure S11). Collectively, these results indicate that the genetic diversity, both at the level of nucleotide and haplotype, of Colombian *Lu. longipalpis* is lower than that of Brazilian populations. Based on this, we hypothesized that the demographic trajectory of *Lu. longipalpis* in Colombia was different from that of the populations in Brazil and thus we examined the trajectory of the effective population size over time of the different lineages of *Lu. longipalpis*. We found that all Brazilian populations experienced an effective population size increase on average around 122,410 generations ago (Figure 6). Captive sandflies have a generation of 6-7 weeks^40^, which would suggest that the population increase occurred between 14,086 and 16,433 years ago (for 6 week or 7 week generations, respectively). While we found the same trend in all Brazilian populations, the historic effective population size of the Colombian population shows no evidence of dramatic change. As we hypothesized, the Colombian and Brazilian populations of *Lu. longipalpis* have undergone different demographic trajectories which may help explain their current levels of genetic diversity.

**Figure 4.**
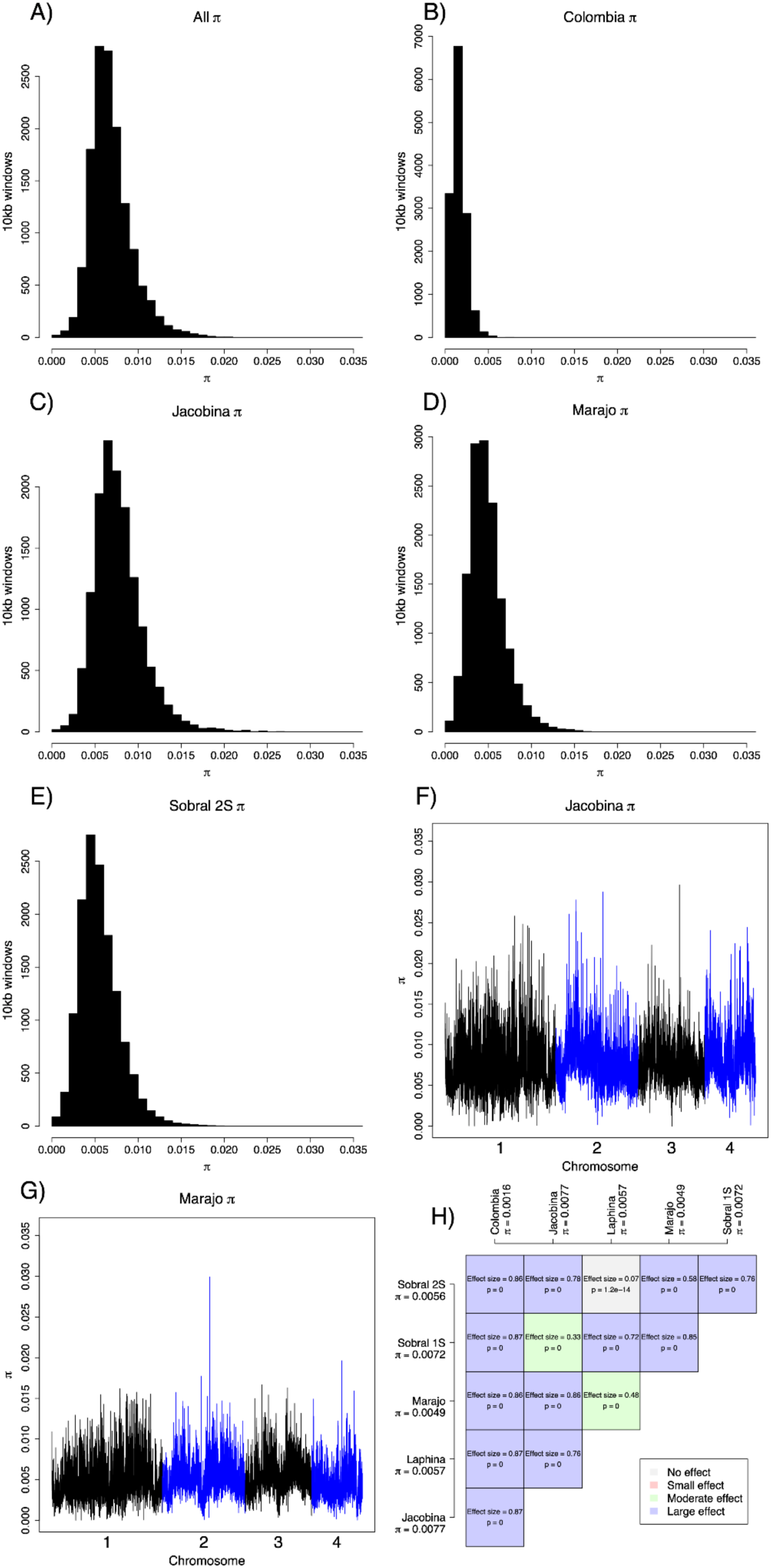
Genetic diversity in different lineages of *Lu. longipalpis*. **A.** Histograms of π in 10kb window sizes along the genome. **B.** π in Colombia **C**. π in Jacobina **D.** π in Marajo **E.** π in Sobral 2S. **F**. π along the genome in Jacobina. **G.** π along the genome in Marajo. Figure 2A and 2B show the π in Colombia. Figure S8 shows π along the genome for the remaining four populations. **H.** Pairwise comparisons show differences in π among pairs of populations.

**Figure 5.**
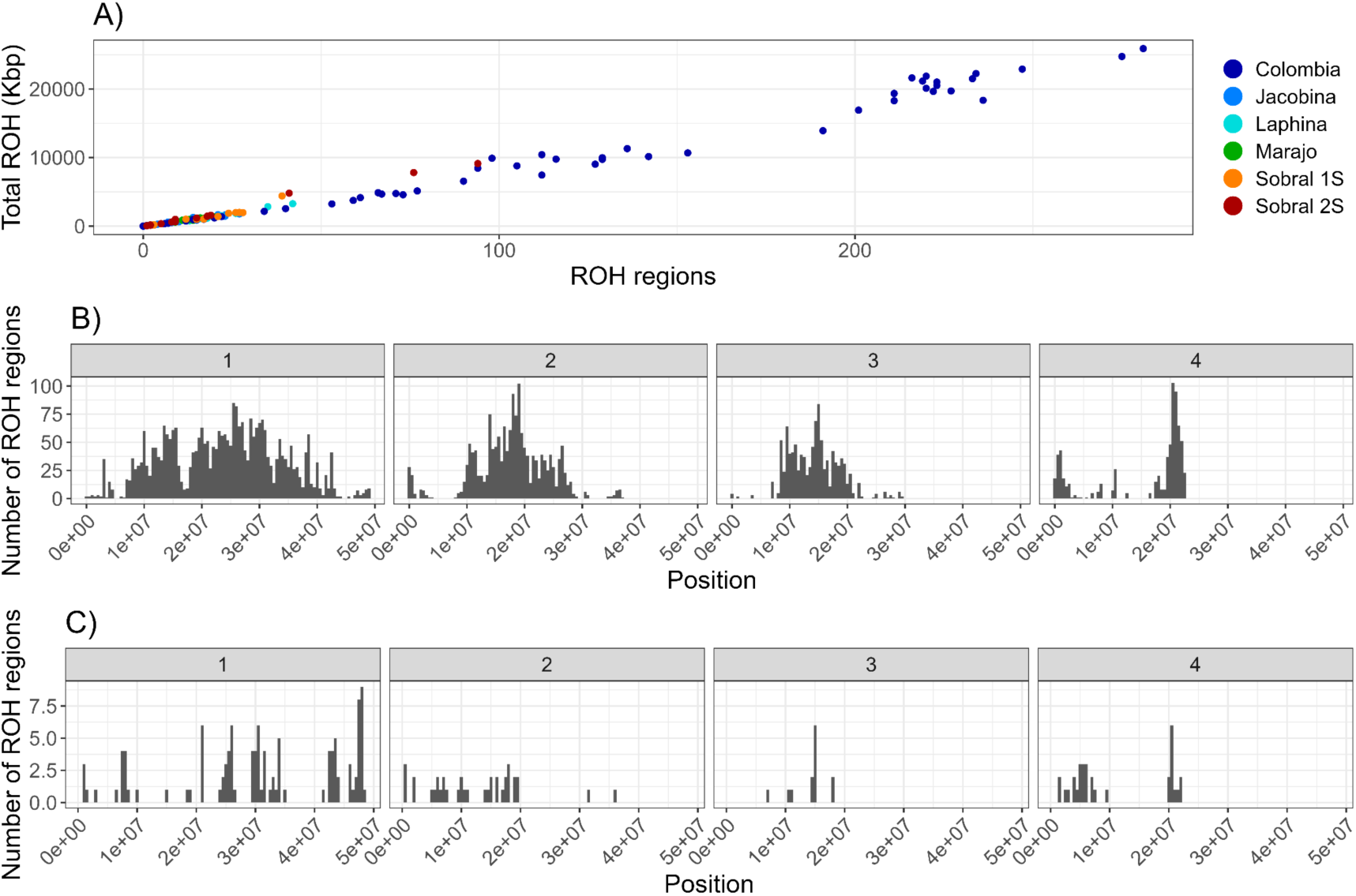
Runs of homozygosity in *Lu. longipalpis*. **A**. Plot depicting the total number of ROH regions over 50kb size and their total length by populations. **B**. Size distribution across genomic regions for Colombian sandflies (binwidth = 500000). **C**. Size distribution across genomic regions for sandflies from Marajo, Brazil (binwidth = 500000). Figure S9 shows the ROH distribution per chromosome for other populations. Figure S10 shows ROH histograms by individual for each population.

**Figure 6.**
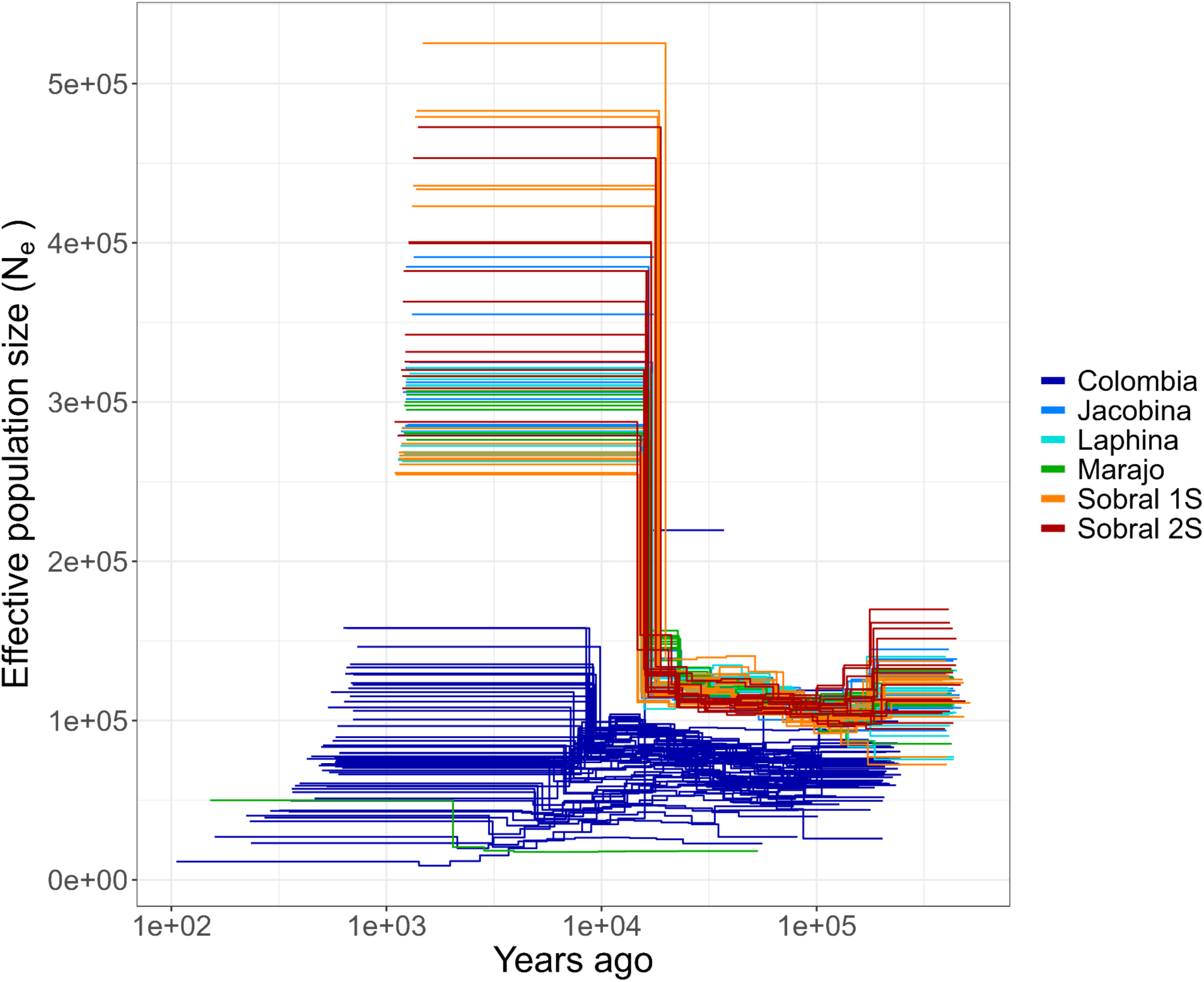
Demographic history of sandflies indicating temporal patterns of predicted effective population size. We assumed a generation time of six weeks.

## DISCUSSION

Phlebotomine sandflies are disease vectors for several human and animal pathogens, and as a result, pose a serious human health threat^41–43^. One of the more notable diseases they transmit is the neglected tropical disease leishmaniasis, and in the Neotropics, one of the most common and important disease vectors of leishmaniasis is *Lutzomyia longipalpis*^44,45^. This sandfly species is widespread across the South American continent^2,14^, poorly characterized^15,16^, and is thought to be increasingly associated with anthropogenic and urbanized habitats^27^. Despite the clear human health risk posed by these vector species, little is known about their basic biology, hampering control and mitigation efforts^46^. Although there are some genomic resources^31,47^, these are limited to Brazilian populations which represent geographically distant populations within a broader cryptic species complex of *Lu. longipalpis*. Here, we present highly continuous genome assemblies for both female and male sandflies from Colombia, which will be a critical resource for future work. We identified signatures of differentiation between the sexes and high divergence across the continent. Further, we discovered that the demographic histories of the different lineages of *Lu. longipalpis sensu lato* are complex, and that genetic information points to the presence of multiple syntopic lineages. We discuss each of these findings in the next paragraphs.

Our first finding is that females and males differ in their genomic composition. Although recent genomic work has indicated no heteromorphy in the Psychodidae^48^, prior work identified the presence of heteromorphic chromosomes in sandflies^49–51^. Here, we sequenced individuals from both sexes and produced highly continuous assemblies of both sexes. We found that the longest chromosome in *Lu. longipalpis* is heteromorphic and ∼1.3 Mbp longer in male sandflies than in female sandflies. Our study builds on these previous observations and reveals that there are differences in TE content and overall genomic composition between the *X* and *Y* chromosomes. Further, our analysis of repeats within the genome indicates that this difference in chromosomal length is likely driven by repeat content in male sandflies, which have an extra ∼1.3 Mbp of repeats in this chromosome. Our results have implications for the study of sex chromosome evolution in dipterans. Further, because only females are hematophagous, and leishmaniasis is known to have negative impacts on longevity and fecundity of sandflies^52,53^, leishmaniasis is likely to be a strong selective force that may impart different selective regimes on female and male genomes, which may be evident either between the sexes or potentially in heteromorphic sex chromosomes.

We identified a large, highly differentiated region between the sexes in a heteromorphic sex chromosome. This region shows a substantial dip in nucleotide diversity in females relative to males and encompasses a suite of genes related to sex determination, including the commonly characterized dipteran sex-determining genes *sex-lethal, fruitless* (*fru*)*, doublesex-and mab-3-related transcription factor A2* (*dsx*)*, lethal(2)*^54^. Some of these genes, *fru* and *dsx*, have been implicated in sex determination in other phlebotomine species^55^. There are also a variety of testes-specific genes (e.g., *ejaculatory bulb-specific protein 3*, *ejaculatory bulb-specific protein 3-like*, *testis-expressed protein 10 homolog*, *motile sperm domain-containing protein*). One gene in this region, *takeout,* shows a high degree of structural conservation and has a conserved role in courtship across insects as diverse as fruit flies, mosquitos, and moths^56^. Other genes in this region of differentiation may be related to basic ecology and life history such as finding mates, finding prey, and effective hematophagy. Among these are several odorant receptor and gustatory receptor genes. Genes in these families play a key role in finding mates and lekking behavior^57^, finding food or blood meals^58–60^, and oviposition site choice^61,62^. In other hematophagous lineages, odorant receptor genes are heavily regulated and exhibit sex-specific and satiation-specific signatures of expression^63^. In *Lu. longipalpis,* pheromones play a key role in attracting mates^18,64–66^; these show population-specific signatures^18^ and are perceived by odorant receptor genes which also show population specific variation ^57^. These may play a role in the cryptic *Lu. longipalpis* species complex via mate choice, and may be a source of prezygotic isolation. Although these roles of odorant receptor genes are well-known, these genes often have disparate functions among dipterans^67^. In sum, there are a suite of genes in the genomic region that are highly differentiated between the sexes that are related to ecology including finding food, finding mates, and oviposition that are potentially playing important, sex-specific roles.

A complementary result, also related to sex differentiation in *Lutzomyia*, is the finding of autosomal alleles associated with sex. In addition to these regions in chromosome 1, our genome wide association study between female and male sandflies in Colombia yielded significant peaks that were intragenic in three predicted genes, one per chromosome on chromosomes 2-4. We were unable to identify the gene product on chromosome 2 via Uniprot, OrthoDB, VectorBase, or NCBI blast. The predicted gene on chromosome 3 has also been predicted in a previous assembly of *Lu. longipalpis* ^31^, but remains an uncharacterized protein product. The intragenic SNPs our GWAS identified as associated with sex differences on chromosome 4 is Apoptotic protease-activating factor 1 (*apaf-1*). The *C. elegans* homolog of this gene (*CED-4*) interacts with the *C. elegans* gene *fem-1* which is crucial for male development^34^ and *apaf-1* deficient mice exhibit male infertility due to degeneration of spermatogonia^35^. The *Drosophila* homolog of *apaf-1* is the gene Apaf-1–related killer (*ark,* or sometimes *dark* for *Drosophila* Apaf-1–related killer), which is required for sperm formation^68^. The SNPs that we identified were present in intronic regions between predicted exons 3 and 4 in our gene annotation. However, the publicly available RNAseq data that we used for annotation only contained female sandflies, potentially missing key, sex-specific isoforms of this gene. We postulate that genomic differentiation between the sexes in this gene is related to sex-specific differences in gametogenesis, or sex conflict (reviewed in ^69^).

Perhaps most notably, our study provides strong evidence of divergence and population structure across the continent. Our analyses combine novel population genomics data for Colombian *Lu. longipalpis* populations with publicly-available whole genome sequence data, which allowed us to carry out the first population genetics study of this critical vector species across South America. Overall divergence between Brazilian and Colombian sandflies was high, supportive of previous work suggesting that *Lu. longipalpis* is a cryptic species complex^15,16,36,37^. In addition, we found evidence that Colombian *Lu. longipalpis* shows lower genetic variability than all of the Brazilian populations. This result manifested in two different facets, lower nucleotide diversity and a substantial increase in ROH tracts. ROH tracts were more common in the *X*-chromosome in all populations but were particularly noticeable in Colombia. It is unclear why Colombian flies exhibit such dramatic patterns of low diversity, particularly because it does not appear to be an isolated population. A potential explanation is reduced vagility in the Colombian lineage. For example if Colombian sandflies roam less during their lifetime than Brazilian flies, they are more likely to experience matings with close relatives and thus show more signs of inbreeding. An additional possibility is that Colombian *Lu. longipalpis* exhibit facultative parthenogenesis, a trait seen in several dipteran species^70^, including in *Lu. mamedei*^71^. Both of these possibilities remain highly speculative and a systematic survey of the reproductive biology of the different lineages of *Lu. longipalpis* is needed to understand the patterns we observed. Our demographic analyses concur with the inference of low genetic diversity in Colombia and indicate that all populations had a similar *Ne* between 16,000 and 14,000 years ago depending on the generation time in the wild; Brazilian populations underwent a large population expansion, Colombian *Lu. longipalpis* did not. This timing is similar to the arrival and expansion of humans in South America^72^. Given that *Lu. longipalpis* is one of the most anthropophilic species, population expansion in Brazilian sandflies may have occurred as a result of human colonization and expansion.

Colombian sandflies (from our single sampling locality) showed structure only between female and male sandflies, whereas multiple Brazilian populations (Jacobina, Sobral) were highly structured in all our analyses. Together, these results indicate the presence of multiple syntopic lineages—a result consistent with the previously hypothesized presence of cryptic species within *Lu. longipalpis*, *sensu lato. Lutzomyia longipalpis* has been proposed to encompass different chemotypes^64^, and in some instances the proposals have been more ambitious and have proposed the existence of a species complex which encompasses different species with stark differences in mating behavior (e.g., mating song, pheromones and lekking behavior). Notably, and despite the differentiation we describe for these lineages, we find some evidence of admixture between lineages. Hybridization seems to be common in *Lutzomyia* (reviewed in^73^), which poses the question of how frequently differentiating lineages exchange alleles. Here we provide evidence for the existence of genome-wide admixture within *Lu. longipalpis*. While this introgression does not appear sufficient to erode the genetic differentiation between lineages, even when they are syntopic, this allele sharing might serve as fodder for adaptation^74,75^. The importance of introgression has been previously hypothesized across the *Lutzomyia* genus^76–78^, and among *Lu. longipalpis* lineages^79,80^; our results represent a formal genome-wide test of the occurrence of this phenomenon in this genus of disease vectors.

Our manuscript leverages the power of whole genome sequencing to understand the evolutionary processes that govern genetic variation in the critical leishmaniasis vector *Lu. longipalpis*. Our results incorporate a genomic perspective into the study of differentiation within *Lu. longipalpis* and represent a resource to not only understand sex chromosome evolution in disease vectors, but also open the door to understanding the molecular and evolutionary mechanisms of the interactions between the vectors, the *Leishmania* parasites, and the hosts affected by the disease. Now that we have a better snapshot of the extent of genetic variation in the vector, the next frontier is understanding whether the evolutionary interactions between sandfly vectors and the *Leishmania* pathogen can provide the key to mitigating the spread of leishmaniasis and the human health threat it poses.

## MATERIALS AND METHODS

### Permits and ethics

All sandflies collections were performed following the guidelines of the Colombian decree No 001566 and 002662. Insects were collected under the permits 001566 and 002662 of the Autoridad de Licencias Ambientales. Sandflies were collected on private property and permission was received from landowners before sampling.

### Sample collection

Samples were collected from a semi-forested location outside of Ricaurte, Cundinamarca, Colombia in November 2021 and October 2022. Sandflies were collected via mechanical aspiration with a mouth aspirator and placed into vials to temporarily hold the insects until processing. Prior to processing, vials containing the sandflies were put in a cooler with ice to immobilize the specimens. Vials were emptied into a Petri dish where the sandflies were selected and classified by species under a Leica dissecting scope. To identify sandfly species, we used taxonomic keys^81,82^. Individual sandflies were removed with tweezers and placed into individual 1.5mL tubes either in liquid nitrogen for genome assembly in 2022 or in RNA-stabilizing buffer (RNALater^TM^; Ambion, USA) for whole genome resequencing in 2021. Samples were exported to North Carolina, USA for sequencing.

### Genome sequencing

Due to their small size (thorax length in females: mean = 0.052 ± 0.0025 mm; males: mean = 0.0483 ± 0.0027 mm;^83^ individuals were pooled prior to genomic DNA extraction for whole genome sequencing (females = 9, males = 10). We used two approaches to obtain reads for the genome assemblies. First, we generated PacBio HiFi reads. We homogenized sandflies with a pestle and incubated them at 55℃ for 30 minutes prior to performing a standard phenol chloroform extraction. Extracted high molecular weight gDNA was sent to the Genomics Core Facility at the Icahn School of Medicine at Mount Sinai, where libraries were prepared using PacBio’s ultra-low DNA input HiFi library prep which utilizes PCR amplification of genomic material. Libraries were pooled and sequenced across two SMRT cells on a Pacific Biosciences Sequel IIe to a coverage of 71.8x and 75.7x coverage for females and males.

Second, we used chromatin configuration. We sent 25 female and 25 male *Lu. longipalpis* in ethanol to Arima Genomics for DNA extraction and sequencing. Hi-C data was generated using the Arima HiC kit and the Arima Library Prep Module according to the Arima Genomics manufacturer’s protocols. Briefly, Arima High Coverage Hi-C is designed to capture the sequence and 3D conformation of genomes. Chromatin from tissues, cell lines, or blood is first crosslinked to preserve genomic structure, then digested with a restriction enzyme cocktail optimized for uniform coverage. The digested ends are labeled with biotinylated nucleotides during a fill-in step and subsequently ligated to capture spatially proximal DNA fragments. These proximally-ligated DNA fragments are purified, fragmented, and enriched for biotin-labeled sequences.The female sample was sequenced to 189.5 million read pairs and the male sample to 180 million read pairs on an Illumina NovaSeq on ‘paired-end’ mode.

### Genome assemblies

We produced separate genome assemblies for the female and male sandflies. While we initially attempted to use HiFiasm^84^, we discovered that haplotype diversity in our pooled data exceeded the allowed diversity in the assembly algorithm. Therefore, we chose to use wtdbg2^85^, an assembler with an algorithm that allows for greater diversity and collapses haplotypes into a single, haploid assembly. We used the “wtpoa-cns” command in wtdbg2 to polish the assembly.

We then scaffolded our initial draft genome assemblies with our Hi-C data. Splits were calculated using Juicer version 1.6^86^ and reads were mapped to the draft assemblies with BWA version 0.7.17^87^. We then ran Samtools version 1.17 to sort the aligned reads and Picard’s “MarkDuplicates” version 2.27.5 to mark duplicates^88^. We indexed our initial contig assembly using Samtools “faidx” prior to running the YaHS (Yet Another Hi-C Scaffolding tool) pipeline^89^ with a MAPQ score of 30. We used Juicer “pre” to create a HiC contact map file which we imported into Juicebox Assembly Tools^90^ to manually curate each assembly. Curation included fixing one major translocation (female assembly) and splitting a chromosome that had an incorrect telomere-to-telomere join (male assembly). We then used Juicer “post” to update the manually curated final assemblies for females and males.

After examination of whole genome sequence results, we identified systemic and dramatic differentiation between female and male sequences along the majority of chromosome 1. We were unable to initially phase a ZW or XY assembly using our assembly approaches, likely due to the pooled nature of our data (i.e., because male sandflies were too small for effective single-fly sequencing, and we were forced to pool multiple individuals). As a result, we took a secondary approach to assemble what we predicted to be a Y chromosome in males using the software SRY^91^ to identify reads that are likely to be from sex-limited chromosomes. We used female and male Illumina whole genome sequence reads to produce sex-specific kmers. We then mapped these kmers to the raw male-only PacBio HiFi reads. We then extracted those reads that were highlighted by presumptive sex-limited kmers from our Illumina data and used wtdbg2^85^ to assemble a Y chromosome from those HiFi reads. We then used RagTag^92^ to scaffold the male Y chromosome using the male X chromosome (chromosome 1) as our backbone. We aligned this to the male “X” chromosome using minimap2^93^ assuming ∼0.1% sequence divergence (“-cx asm5”) and visualized the alignment using “*pafr*” in R ^94^.

After producing our scaffolded assemblies, we identified repeat elements in the assemblies. We used Repeat Modeler version 2.0.4^95^ to model repeat elements within each assembly. Most of these repeat elements were unidentified by Repeat Modeler, so we ran Transposon Classifier RFS^96^ to classify these repeat regions. We then created a dipteran and *Lu. longipalpis* specific database of repeat elements. We combined the curated DFAM database version 3.4^97^ with the RepBase version 25.05 within RepeatMasker version 4.1.2-p1^98^. We used the function “famdb.py” to extract all dipteran sequences from these databases, then added our sex-specific sandfly repeat databases. We then identified and soft-masked repeats throughout the genome using RepeatMasker version 4.1.2-p1^98^ and specified our curated vertebrate and *Lu. longipalpis* repeat library as the library of repeats.

We then annotated assemblies using Braker version 3.0.8^99–102^. We used a combination of protein evidence and RNA seq evidence to annotate putative protein coding regions of the genome. We used the OrthoDB v11 database^103^ and the VectorBase 67 database^104,105^ as input protein data for annotation. Protein sequences were aligned to the genomes using miniprot^106^. We used the SRA toolkit^107^ to download publicly-available RNA seq data from *Lu. longipalpis* to annotate the genome. We downloaded samples with accession numbers SRR535756-SRR535778. RNA seq reads were mapped to our genomes using HISAT2^108^ and sorted/manipulated using samtools^109^. We followed this with Braker3, utilizing the GeneMark-ETP pipeline^110^ to predict protein coding genes from RNAseq evidence. This was then compared to the protein evidence, OrthoDB and VectorBase in this case, to find homologous proteins that were then mapped back to the genome with ProtHint^101^. Augustus^111,112^ then predicted a second set of genes. We additionally used the diptera_odb10 database^113^ in Augustus to predict potentially missing protein products. We then used TSEBRA^114^ to merge gene predictions from Augustus and the GeneMark-ETP pipelines.

### WGS resequencing

We resequenced a total of 63 *Lu. longipalpis* from Ricaurte, Cundinamarca, Colombia. Of these, 37 sandflies were female and 26 were males sample information in Table S1). Briefly, we conducted a homogenization and incubation similar to that of the WGS gDNA extraction. Following the incubation, we followed a modified version of the instructions on the Zymo Genomic DNA Clean and Concentrator kit, and quantified eluted DNA using a Qubit. Our full extraction protocol can be found in the supplemental materials. Library prep and sequencing were conducted at the University of North Carolina-Chapel Hill’s High-Throughput Sequencing Facility. Libraries were prepped using a ThruPLEX method for low DNA concentrations (CHiP-seq). Following library preparation, pooled paired-end 150bp reads were sequenced on an Illumina NovaSeq 6000 SP.

### WGS mapping, variant calling, and diversity summary statistics

Next we measure the extent of genetic diversity and differentiation within *Lu. longipalpis*. We followed the GATK best practices for variant calling^115^. We used BBMap version 38.87 (https://github.com/BioInfoTools/BBMap) to trim adapters from whole genome resequencing data and then mapped reads to the female reference genome using BWA version 0.7.17^87^. Read groups were added using GATK’s AddOrReplaceReadGroups. For samples sequenced across two lanes, aligned data were then merged using the “merge” tool in samtools version 1.14^109^. Aligned data were cleaned with Picard’s CleanSam, sorted with SortSam, and duplicates marked with MarkDuplicates (Picard version 2.23.4^88^. Genotypes were then called with GATK HaplotypeCaller, before joint genotyping data with GenotypeGVCFs. All GATK functions were from GATK version 4.1.9.0.

We merged bam files for each sample that were sequenced on multiple lanes using samtools version 1.14 merge function^109^. We generated a Beagle file of genotype probabilities using *angsd* version: 0.921-11-g20b0655, and then calculated principal components using the *angsd* script “pcangsd.py”^116^. We then extracted eigenvalues and calculated the percent of the variance in our data explained by each PC axis, which we then plotted in R version 4.1.1 (“Kick Things”) using ggplot2^117^. In addition to running this PCA pipeline with whole genome data, we subsetted the alignment bam file in order to separately examine differentiation between the presumptive sex chromosome (chromosome 1) and the autosomes (chromosomes 2-4) using the same approach.

We examined basic population statistics (nucleotide diversity, π, and population divergence [F_ST_, Dxy]) using pixy^118^, which accounts for invariant sites. We calculated pairwise differences in nucleotide diversity and Weir-Cockerham F_ST_ values^119^. We were particularly interested in potential differentiation between female and male sandflies, given that only females are hematophagous and thus only females are vectors of leishmaniasis. Per pixy’s instructions, we first created a vcf file of invariant sites using the “-all-sites” flag in GATK’s GenotypeGVCFs. We then filtered the corresponding gvcf file to remove sites with a high proportion of missing data (--max-missing 0.9), low coverage (--min-meanDP 3), and high overall depth (--max-meanDP 40) using vcftools 0.1.15^120^. We then used pixy to calculate F_ST_ and Dxy across 1,000bp windows between female and male sandflies. The window size was chosen five the extent of LD in *Lu. longipalpis*.) . Additionally, we subset our gvcf file into a female-only and a male-only gvcf which we used to calculate nucleotide diversity (π) across 1,000bp windows in female and male sandflies. We used PLINK 1.90b6.21^121^ to calculate linkage disequilibrium. We additionally calculated Tajima’s D in 1,000 bp windows using VCFtools.

We detected runs of homozygosity (ROH) using PLINK 1.90b6.21^121^ with the parameters: --allow-extra-chr and --homozyg-kb 5. This command estimates the run length and identifies contiguous segments of homozygosity greater than 5 kbp by assessing the observed genotype data against expected patterns of homozygosity. ROH was detected separately for each of Colombia and the five Brazil populations. With a 5 kbp minimum size, we identified a minimum of 100 SNPs per region and on average there were 189-276 SNPs per region. We repeated the same analyses using different parameter values for the minimum length of ROH in kbp (5, 10, 25, 50, 100), and results were similar despite fewer ROH regions with higher kb thresholds.

### Phylogenetic trees

We used trees to address two questions. First, to examine the genealogy of the largest chromosome relative to the presumptive autosomes in placement of females and males in relation to each other, and second, to examine continent-wide evolutionary relationships. We used the python script “vcf2phylp.py”^122^ to split our vcf files into 100,000 bp regions and convert them into the phylip format.

To examine the genealogy of the largest chromosome relative to the presumptive autosomes, we created two separate trees using 1) only the largest chromosome or 2) the presumptive autosomes in just Colombian sandflies. For both trees, we made a phylogenetic tree for each window using IQ-TREE^123^ version 2.2.2.7 with DNA specified (“-st DNA”) and 1000 bootstraps (“-bb 1000”). We merged the resulting consensus tree files into a single tree and used ASTRAL^124^ version 5.7.8 to create a single genome-wide consensus tree. We additionally created trees solely from chromosome 1 and the autosomes.

To examine evolutionary relationships across the continent, we made a single genome-wide tree using the above methods. The approach for this question is identical to the one described immediately above, with the exception that we created a single tree including all chromosomes. Further, we examined gene concordance between the genome wide tree and conserved, core dipteran genes from the BUSCO diptera_odb10 database^113^. To do this, we used ASTRAL^124^ version 5.7.8 to create a single genome-wide consensus tree that is constrained to the populations sampled. We additionally extracted all the complete, single copy dipteran BUSCO genes assembled in our male assembly, using the python script “vcf2phylp.py”^122^ to convert the vcf of these genomic regions into the phylip format. We created single gene trees as per the windows above using IQ-TREE, and created a single consensus population-level tree using ASTRAL as we did for our whole genome tree. We then used IQ-TREE^123^ version 2.2.2.7 to calculate gene concordance between each gene tree and the overall consensus tree. Finally, we examined gene flow and the potential for introgression using fbranch statistics in Dsuite ^125^. Briefly, we used our consensus population tree and our variant call file as input to examine fbranch statistics using “Dtrios” to examine all trios and “Fbranch” to examine the output fbranch statistics. Given the clear divergence between Brazilian and Colombian sandflies, we used Colombia as the outgroup in our analysis and focused on Brazilian sandflies.

### Population genomics of geographically disparate *Lu. longipalpis* populations

*Lutzomyia longipalpis* has a broad geographic range from Argentina through Mexico^2,14^. Further, multilocus sequence typing and phenotypic surveys have revealed strong population structure which might correspond to cryptic speciation^17,18^ and divergences in courtship song^20^, morphology,^21^, and pheromones^18^. Recent work sequenced Brazilian *Lu. longipalpis* from four localities in Brazil^31^. Here we examined basic population genomic metrics between Colombian and Brazilian *Lu. longipalpis.* We were particularly interested in differentiation between populations of putative *Lu. longipalpis,* and as a result focus on metrics such as F_ST_ and Dxy between populations.

Out of the 68 Colombian samples, 63 sandflies (37 females and 26 males) passed our quality metrics for population level analyses (>50% of reads mapping to our *Lu. longipalpis* female assembly and > 5X average coverage). On average, 91.4% of female reads and 84.4% of male reads mapped to the female assembly. Average coverage was 16.8x and 13.2x for females and males respectively. When mapped to the male assembly, 84.3% of female reads and 79.6% of male reads mapped and we had 14.6x and 11.7x coverage for females and males respectively.

Additionally, we downloaded whole genome resequence data from^31^ using the SRA toolkit^107^. Table S1 lists the accession numbers of the previously published reads used in this study. We then followed the same protocol that we used for our Colombian samples, the GATK best-practices for variant calling ^115^. We used BBMap version 38.87 (https://github.com/BioInfoTools/BBMap) to trim adaptors from whole genome resequencing data and then mapped reads to the female reference genome using BWA version 0.7.17^87^. Read groups were added using GATK’s AddOrReplaceReadGroups. For samples sequenced across two lanes, aligned data were then merged using the “merge” tool in samtools version 1.14^109^. Aligned data were cleaned with Picard’s CleanSam, sorted with SortSam, and duplicates marked with MarkDuplicates (Picard version 2.23.4^88^. Genotypes were then called with GATK HaplotypeCaller, before joint genotyping data with GenotypeGVCFs. All GATK functions were from GATK version 4.1.9.0.

For samples across the continent of South America, we examined summary population statistics (π, F_st_, and Dxy) as we did for samples within Colombia (see above). We filtered our gvcf file to remove sites with a high proportion of missing data (--max-missing 0.9), low coverage ( --min-meanDP 3), and high overall depth (--max-meanDP 40) using vcftools 0.1.15^120^. We then used pixy to calculate F_ST_ and Dxy across 1,000bp windows between populations. Additionally, we subset our gvcf file into a female-only and a male-only gvcf which we used to calculate nucleotide diversity (π) across 1,000bp windows in female and male sandflies. We additionally calculated Tajima’s D in 1,000 bp windows using VCFtools.

In order to examine population structure across the continent, we used angsd^116^ to examine the data with a PCA, structure, and *ADMIXTURE* based approaches. We used a similar approach to the within-Colombia examinations. We generated a Beagle file of genotype probabilities using *angsd* version: 0.921-11-g20b0655, and then calculated principal components and genotype probabilities using the *angsd* script “pcangsd.py”^116^. This program calculates ancestry based on individual allele frequency and uses a similar approach to delta K in that it permutes iteratively across a range of ancestry possibilities. We then extracted eigenvalues and calculated the percent variance in our data explained and plotted both this PCA and a STRUCTURE plot in R version 4.1.1 (“Kick Things”) using ggplot2^117^.

To quantify if sex was responsible for differences in genome size, we used a linear model (function “lm” in R) with average π per window in the sexes as the outcome and sex, chromosome, and the interaction of sex and chromosome as fixed effects.

We estimated demographic histories using MSMC2^126^ with one thread and ten time segments of different lengths using the parameters ‘-t 1 -p 1-6+8-1+1-4’. MSMC2 was run separately for individual samples since we had unphased genotypes. The MSCM2 input files were generated using msmc-tools^127^. Separate input files were generated for each of the four major chromosomes using ‘generate_multihetsep.py’ and then concatenated together into separate genomic input files for each individual sample. ‘generate_multihetsep.py’ requires two input files: a mask of sites with coverage for that individual and a genomic mappability mask. The covered site masks were created in three steps run separately for each chromosome and each individual sample using Samtools 1.21^128^, BCFtools^128^ and msmc-tools. The genomic mappability mask was created using genmap^129^. MSMC2 results were plotted using R ^130^ using a mutation rate of *μ*=1.25e-8 and an estimated generation time of 49 days^40^ with time in years calculated as ‘time_boundary / *μ* / gen’. The effective population size was calculated as (1 / **λ**) / (2 * *μ*) where lambda is the coalescence rate returned by MSMC2.

### Genome wide association study for sex determination

Finally, we ran a genome-wide association study (GWAS) for 133 samples from both Colombia and Brazil using sex as the phenotype (113 females, 20 males) to identify genetic regions associated with sex determination. First, we created the required plink input files from our vcf by filtering for just the four chromosomes and using a minor allele frequency of 0.05. To run the GWAS, we used plink 2.0^121^ using a generalized linear model framework with sex as a binary variable^131^, and we filtered out SNPs with 20% or more missing data (--geno 0.2) and a minor allele frequency of 0.05 (--maf 0.05). Results are presented from the continent wide analysis, but results are the same using just the Colombian samples yielded qualitatively identical results.

## Data availability

All raw sequence data as well as genome assemblies are available on NCBI under BioProject PRJNA1110158. Code can be found at https://github.com/AdamStuckert/Lu_longipalpis; and will be deposited in Zenodo upon acceptance.

## Author contributions

AMMS: Conceived project, collected samples, conducted wet lab work, analyzed data, wrote manuscript

DT: Analyzed data, wrote manuscript

ERRD: Conducted wet lab work, wrote manuscript

DP: Conducted wet lab work, wrote manuscript

MP: Conducted wet lab work, wrote manuscript

CM: Conceived project

JAR: Wrote manuscript

EB: Conceived project, collected samples

JRW: Conducted wet lab work, wrote manuscript

RV: Conceived project, collected samples, identified sandflies, wrote manuscript

DRM: Conceived project, collected samples, analyzed data, wrote manuscript

## Acknowledgments

The research presented here was supported by the National Institute of General Medical Sciences under award R35GM148244.

