## Supplement for "Sex differentiation, divergence, and introgression in the leishmaniasis vector *Lutzomyia longipalpis*"

**SUPPLEMENTARY TABLES**

Table S1. Sample information for short read genomic data used in this study. This includes the collections from Colombia and a list of accession numbers of the previously published reads used in this study.

| Run | Site | Sex |
|---|---|---|
| LU_LONG_M_131_CAATTAAC-CGAGATAT_S34 | Ricaurte, Colombia | Male |
| JB_LM30-lutzomyia_male_42_021921_TCCTGAGC-AAGGCTAT_S79 | Ricaurte, Colombia | Male |
| JB_LM11-lutzomyia_male_23_021921_CGTACTAG-CCTAGAGT_S58 | Ricaurte, Colombia | Male |
| JB_LF33-lutzomyia_female_100_021921_CGAGGCTG-GAGCCTTA_S26 | Ricaurte, Colombia | Female |
| JB_LF13-lutzomyia_female_80_021921_TAGGCATG-CCTAGAGT_S4 | Ricaurte, Colombia | Female |
| JB_LF4-lutzomyia_female_71_021921_GGACTCCT-TTCTAGCT_S44 | Ricaurte, Colombia | Female |
| JB_LM3-lutzomyia_male_15_021921_TAAGGCGA-CCTAGAGT_S80 | Ricaurte, Colombia | Male |
| LU_LONG_F_P12_GGACTTGG-CGCAGACG_S20 | Ricaurte, Colombia | Female |
| JB_LM1-lutzomyia_male_13_021921_TAAGGCGA-TCGACTAG_S67 | Ricaurte, Colombia | Male |
| JB_LF52-lutzomyia_female_119_021921_GCTCATGA-TTCTAGCT_S47 | Ricaurte, Colombia | Female |
| JB_LM24-lutzomyia_male_36_021921_AGGCAGAA-TTATGCGA_S72 | Ricaurte, Colombia | Male |
| JB_LM22-lutzomyia_male_34_021921_AGGCAGAA-AAGGCTAT_S70 | Ricaurte, Colombia | Male |
| JB_LM19-lutzomyia_male_31_021921_AGGCAGAA-CCTAGAGT_S66 | Ricaurte, Colombia | Male |
| JB_LM5-lutzomyia_male_17_021921_TAAGGCGA-CTATTAAG_S82 | Ricaurte, Colombia | Male |
| JB_LF45-lutzomyia_female_112_021921_GTAGAGGA-CCTAGAGT_S39 | Ricaurte, Colombia | Female |
| JB_LF51-lutzomyia_female_118_021921_GCTCATGA-TCGACTAG_S46 | Ricaurte, Colombia | Female |

| | | |
|---|---|---|
| JB_LM18-lutzomyia_male_30_021921_AGGCAGAA-TTCTAGCT_S65 | Ricaurte, Colombia | Male |
| JB_LM2-lutzomyia_male_14_021921_TAAGGCGA-TTCTAGCT_S78 | Ricaurte, Colombia | Male |
| LU_LONG_F_138_CATAGAGT-TGCCACCA_S5 | Ricaurte, Colombia | Female |
| JB_LF48-lutzomyia_female_115_021921_GTAGAGGA-AAGGCTAT_S42 | Ricaurte, Colombia | Female |
| LU_LONG_F_147_CTCACCAA-CTAGGCAA_S14 | Ricaurte, Colombia | Female |
| JB_LF22-lutzomyia_female_89_021921_CTCTCTAC-GCGTAAGA_S14 | Ricaurte, Colombia | Female |
| JB_LM7-lutzomyia_male_19_021921_TAAGGCGA-GAGCCTTA_S84 | Ricaurte, Colombia | Male |
| LU_LONG_M_127_GCCACAGG-CATGCCAT_S30 | Ricaurte, Colombia | Male |
| JB_LF7-lutzomyia_female_74_021921_GGACTCCT-CTATTAAG_S54 | Ricaurte, Colombia | Female |
| LU_LONG_F_P19_AGTTCAGG-CCAACAGA_S26 | Ricaurte, Colombia | Female |
| JB_LF18-lutzomyia_female_85_021921_TAGGCATG-TTATGCGA_S9 | Ricaurte, Colombia | Female |
| LU_LONG_F_154_GTATGTTC-AACAGGAA_S18 | Ricaurte, Colombia | Female |
| LU_LONG_M_128_ATTGTGAA-TGCATTGC_S31 | Ricaurte, Colombia | Male |
| LU_LONG_F_135_GACGTCTT-GGTTCACC_S3 | Ricaurte, Colombia | Female |
| JB_LM10-lutzomyia_male_22_021921_CGTACTAG-TTCTAGCT_S57 | Ricaurte, Colombia | Male |
| LU_LONG_F_150_CGCTATGT-TCCGACAC_S17 | Ricaurte, Colombia | Female |
| LU_LONG_F_P11_TTATAACC-GATATCGA_S19 | Ricaurte, Colombia | Female |
| JB_LF11-lutzomyia_female_78_021921_TAGGCATG-TCGACTAG_S2 | Ricaurte, Colombia | Female |
| LU_LONG_F_146_GAACCGCG-TGACCTTA_S13 | Ricaurte, Colombia | Female |

| | | |
|---|---|---|
| JB_LF40-lutzomyia_female_107_021921_AAGAGGCA-AAGGCTAT_S34 | Ricaurte, Colombia | Female |
| LU_LONG_F_P18_TGGATCGA-GTGCGATA_S25 | Ricaurte, Colombia | Female |
| LU_LONG_F_137_TGCGAGAC-CATTGTTG_S4 | Ricaurte, Colombia | Female |
| JB_LF25-lutzomyia_female_92_021921_CTCTCTAC-GAGCCTTA_S17 | Ricaurte, Colombia | Female |
| LU_LONG_F_143_CTGCTTCC-GATAGATC_S10 | Ricaurte, Colombia | Female |
| JB_LF47-lutzomyia_female_114_021921_GTAGAGGA-CTATTAAG_S41 | Ricaurte, Colombia | Female |
| JB_LM17-lutzomyia_male_29_021921_AGGCAGAA-TCGACTAG_S64 | Ricaurte, Colombia | Male |
| LU_LONG_F_P16_GCTTGTCA-GAACATAC_S23 | Ricaurte, Colombia | Female |
| JB_LF43-lutzomyia_female_110_021921_GTAGAGGA-TCGACTAG_S37 | Ricaurte, Colombia | Female |
| LU_LONG_F_149_TATCGCAC-CTTAGTGT_S16 | Ricaurte, Colombia | Female |
| JB_LF5-lutzomyia_female_72_021921_GGACTCCT-CCTAGAGT_S52 | Ricaurte, Colombia | Female |
| LU_LONG_F_144_TCATCCTT-AGCGAGCT_S11 | Ricaurte, Colombia | Female |
| JB_LM25-lutzomyia_male_37_021921_TCCTGAGC-TCGACTAG_S73 | Ricaurte, Colombia | Male |
| LU_LONG_F_142_GGTCACGA-GTATTATG_S9 | Ricaurte, Colombia | Female |
| LU_LONG_F_145_AGGTTATA-CAGTTCCG_S12 | Ricaurte, Colombia | Female |
| LU_LONG_F_P20_GACCTGAA-TTGGTGAG_S27 | Ricaurte, Colombia | Female |
| JB_LM16-lutzomyia_male_28_021921_CGTACTAG-TTATGCGA_S63 | Ricaurte, Colombia | Male |
| JB_LM14-lutzomyia_male_26_021921_CGTACTAG-AAGGCTAT_S61 | Ricaurte, Colombia | Male |
| LU_LONG_F_140_GTGAATAT-TCTCATTC_S7 | Ricaurte, Colombia | Female |

| | | |
|---|---|---|
| LU_LONG_F_148_TCTGTTGG-TCGAATGG_S15 | Ricaurte, Colombia | Female |
| LU_LONG_F_141_AACTGTAG-ACGCCGCA_S8 | Ricaurte, Colombia | Female |
| LU_LONG_F_P14_AAGTCCAA-TATGAGTA_S21 | Ricaurte, Colombia | Female |
| JB_LM13-lutzomyia_male_25_021921_CGTACTAG-CTATTAAG_S60 | Ricaurte, Colombia | Male |
| LU_LONG_F_P17_CAAGCTAG-ACATAGCG_S24 | Ricaurte, Colombia | Female |
| LU_LONG_F_139_ACAGGCGC-CTCTGCCT_S6 | Ricaurte, Colombia | Female |
| LU_LONG_F_P15_ATCCACTG-AGGTGCGT_S22 | Ricaurte, Colombia | Female |
| JB_LF26-lutzomyia_female_93_021921_CTCTCTAC-TTATGCGA_S18 | Ricaurte, Colombia | Female |
| JB_LF15-lutzomyia_female_82_021921_TAGGCATG-CTATTAAG_S6 | Ricaurte, Colombia | Female |
| SRR3109093 | Jacobina, Brazil | Female |
| SRR3109094 | Jacobina, Brazil | Female |
| SRR3109095 | Jacobina, Brazil | Female |
| SRR3109096 | Jacobina, Brazil | Female |
| SRR3109097 | Jacobina, Brazil | Female |
| SRR3109098 | Jacobina, Brazil | Female |
| SRR3109099 | Jacobina, Brazil | Female |
| SRR3109100 | Jacobina, Brazil | Female |
| SRR3109101 | Jacobina, Brazil | Female |
| SRR3109102 | Jacobina, Brazil | Female |
| SRR3109103 | Jacobina, Brazil | Female |
| SRR3109104 | Jacobina, Brazil | Female |
| SRR3109105 | Jacobina, Brazil | Female |
| SRR3109106 | Jacobina, Brazil | Female |
| SRR3109107 | Laphina, Brazil | Female |
| SRR3109108 | Laphina, Brazil | Female |

| | | |
|---|---|---|
| SRR3109109 | Laphina, Brazil | Female |
| SRR3109110 | Laphina, Brazil | Female |
| SRR3109111 | Laphina, Brazil | Female |
| SRR3109112 | Laphina, Brazil | Female |
| SRR3109113 | Laphina, Brazil | Female |
| SRR3109114 | Laphina, Brazil | Female |
| SRR3109115 | Laphina, Brazil | Female |
| SRR3109116 | Laphina, Brazil | Female |
| SRR3109117 | Laphina, Brazil | Female |
| SRR3109118 | Laphina, Brazil | Female |
| SRR3109119 | Laphina, Brazil | Female |
| SRR3109120 | Laphina, Brazil | Female |
| SRR3109121 | Laphina, Brazil | Female |
| SRR3109122 | Marajo, Brazil | Female |
| SRR3109123 | Marajo, Brazil | Female |
| SRR3109124 | Marajo, Brazil | Female |
| SRR3109125 | Marajo, Brazil | Female |
| SRR3109126 | Marajo, Brazil | Female |
| SRR3109127 | Marajo, Brazil | Female |
| SRR3109128 | Marajo, Brazil | Female |
| SRR3109129 | Marajo, Brazil | Female |
| SRR3109130 | Marajo, Brazil | Female |
| SRR3109131 | Marajo, Brazil | Female |
| SRR3109132 | Sobral 1S, Brazil | Female |
| SRR3109133 | Sobral 1S, Brazil | Female |
| SRR3109134 | Sobral 1S, Brazil | Female |
| SRR3109135 | Sobral 1S, Brazil | Female |

| | | |
|---|---|---|
| SRR3109136 | Sobral 1S, Brazil | Female |
| SRR3109137 | Sobral 1S, Brazil | Female |
| SRR3109138 | Sobral 1S, Brazil | Female |
| SRR3109139 | Sobral 1S, Brazil | Female |
| SRR3109140 | Sobral 1S, Brazil | Female |
| SRR3109141 | Sobral 1S, Brazil | Female |
| SRR3109142 | Sobral 1S, Brazil | Female |
| SRR3109143 | Sobral 1S, Brazil | Female |
| SRR3109144 | Sobral 1S, Brazil | Female |
| SRR3109145 | Sobral 1S, Brazil | Female |
| SRR3109146 | Sobral 1S, Brazil | Female |
| SRR3109148 | Sobral 2S, Brazil | Female |
| SRR3109149 | Sobral 2S, Brazil | Female |
| SRR3109150 | Sobral 2S, Brazil | Female |
| SRR3109151 | Sobral 2S, Brazil | Female |
| SRR3109152 | Sobral 2S, Brazil | Female |
| SRR3109153 | Sobral 2S, Brazil | Female |
| SRR3109154 | Sobral 2S, Brazil | Female |
| SRR3109155 | Sobral 2S, Brazil | Female |

| | | |
|---|---|---|
| SRR3109156 | Sobral 2S, Brazil | Female |
| SRR3109157 | Sobral 2S, Brazil | Female |
| SRR3109158 | Sobral 2S, Brazil | Female |
| SRR3109159 | Sobral 2S, Brazil | Female |
| SRR3109160 | Sobral 2S, Brazil | Female |
| SRR3109161 | Sobral 2S, Brazil | Female |
| SRR3109162 | Sobral 2S, Brazil | Female |
| SRR3109163 | Sobral 2S, Brazil | Female |

Table S2. Significant GWAS hits. The table shows the name, location, association p-value, and known orthologs in other dipterans.

| chrom | pos | p | gene_type | gene_id | gene_distance | Flybase_gene_name | gene_name |
|---|---|---|---|---|---|---|---|
| 2 | 11287131 | 1.82E-08 | Downstream | g4778 | 2049 | | |
| 2 | 11287131 | 1.82E-08 | Upstream | g4777 | 23544 | | |
| 2 | 17800961 | 5.17E-09 | Downstream | g5201 | 165 | | uncharacterized protein |
| 2 | 17800961 | 5.17E-09 | Upstream | g5200 | 35253 | | eye-specific diacylglycerol kinase-like |
| 2 | 19878643 | 1.12E-09 | Genic | g5391 | | | |
| 3 | 3650443 | 2.59E-09 | Downstream | g6882 | 36835 | CG17870 | 14-3-3 protein zeta |
| 3 | 3650443 | 2.59E-09 | Upstream | g6881 | 1646 | | |
| 3 | 8753610 | 8.95E-08 | Genic | g7272 | | CAA65152.1 | mdg3 element ORF |

| | | | | | | | |
|---|---|---|---|---|---|---|---|
| 3 | 8756972 | 4.43 E-09 | Genic | g7272 | | CAA65152.1 | mdg3 element ORF |
| 3 | 8757227 | 5.06 E-08 | Genic | g7272 | | CAA65152.1 | mdg3 element ORF |
| 3 | 14121068 | 2.91 E-09 | Downstream | g7711 | 14311 | | |
| 3 | 14121068 | 2.91 E-09 | Upstream | g7710 | 14987 | | |
| 3 | 14909442 | 1.22 E-08 | Downstream | g7789 | 4284 | | Orf2 |
| 3 | 14909442 | 1.22 E-08 | Upstream | g7788 | 6504 | CG15715 | zinc finger protein 706 |
| 3 | 14909452 | 4.62 E-09 | Downstream | g7789 | 4274 | | Orf2 |
| 3 | 14909452 | 4.62 E-09 | Upstream | g7788 | 6514 | CG15715 | zinc finger protein 706 |
| 3 | 15059348 | 4.94 E-08 | Downstream | g7799 | 3787 | | integrator complex subunit 12 #(low query coverage/confidence) |
| 3 | 15059348 | 4.94 E-08 | Upstream | g7798 | 8033 | | uncharacterized protein K02A2.6-like |
| 3 | 15061297 | 2.94 E-08 | Downstream | g7799 | 1838 | | integrator complex subunit 12 #(low query coverage/confidence) |
| 3 | 15061297 | 2.94 E-08 | Upstream | g7798 | 9982 | | uncharacterized protein K02A2.6-like |
| 3 | 15062377 | 2.89 E-09 | Downstream | g7799 | 758 | | integrator complex subunit 12 #(low query coverage/confidence) |
| 3 | 15062377 | 2.89 E-09 | Upstream | g7798 | 11062 | | uncharacterized protein K02A2.6-like |
| 3 | 27024829 | 7.38 E-09 | Downstream | g8767 | 7639 | AAD37797 | pleiotrophin isoform |
| 3 | 27024829 | 7.38 E-09 | Upstream | g8766 | 15256 | | |
| 3 | 27024841 | 3.28 E-08 | Downstream | g8767 | 7627 | AAD37797 | pleiotrophin isoform |

| | | | | | | | |
|---|---|---|---|---|---|---|---|
| 3 | 27024841 | 3.28E-08 | Upstream | g8766 | 15268 | | |
| 3 | 27024846 | 4.65E-08 | Downstream | g8767 | 7622 | AAD37797 | pleiotrophin isoform |
| 3 | 27024846 | 4.65E-08 | Upstream | g8766 | 15273 | | |
| 3 | 27024848 | 4.27E-08 | Downstream | g8767 | 7620 | AAD37797 | pleiotrophin isoform |
| 3 | 27024848 | 4.27E-08 | Upstream | g8766 | 15275 | | |
| 4 | 89619 | 6.71E-08 | Downstream | g8996 | 39230 | | na |
| 4 | 89619 | 6.71E-08 | Upstream | g8995 | 10970 | | LIM/homeobox protein Lhx2 #(low confidence) |
| 4 | 477670 | 1.12E-09 | Genic | g9046 | | CG6829 | uncharacterized protein |
| 4 | 477673 | 1.12E-09 | Genic | g9046 | | CG6829 | uncharacterized protein |
| 4 | 477798 | 1.03E-08 | Genic | g9046 | | CG6829 | uncharacterized protein |
| 4 | 477808 | 3.89E-09 | Genic | g9046 | | CG6829 | uncharacterized protein |
| 4 | 4318823 | 1.76E-09 | Downstream | g9424 | 853 | CG2095 | exocyst complex component 4 |
| 4 | 4318823 | 1.76E-09 | Upstream | g9423 | 355 | AAL13990.1 | heat shock factor protein |
| 4 | 16129633 | 1.33E-09 | Downstream | g10362 | 4748 | CG8333 | enhancer of split mgamma protein |
| 4 | 16129633 | 1.33E-09 | Upstream | g10361 | 18627 | | |
| 4 | 16129745 | 1.13E-09 | Downstream | g10362 | 4636 | CG8333 | enhancer of split mgamma protein |
| 4 | 16129745 | 1.13E-09 | Upstream | g10361 | 18739 | | |
| 4 | 16129961 | 7.69E-08 | Downstream | g10362 | 4420 | CG8333 | enhancer of split mgamma protein |
| 4 | 16129961 | 7.69E-08 | Upstream | g10361 | 18955 | | |

| | | | | | | | |
|---|---|---|---|---|---|---|---|
| 4 | 16130036 | 1.61E-09 | Downstream | g10362 | 4345 | CG8333 | enhancer of split mgamma protein |
| 4 | 16130036 | 1.61E-09 | Upstream | g10361 | 19030 | | |
| 4 | 16130047 | 1.61E-09 | Downstream | g10362 | 4334 | CG8333 | enhancer of split mgamma protein |
| 4 | 16130047 | 1.61E-09 | Upstream | g10361 | 19041 | | |
| 4 | 16131132 | 1.82E-08 | Downstream | g10362 | 3249 | CG8333 | enhancer of split mgamma protein |
| 4 | 16131132 | 1.82E-08 | Upstream | g10361 | 20126 | | |
| 4 | 18327554 | 9.57E-09 | Downstream | g10493 | 670 | CG6975 | tuberin |
| 4 | 18327554 | 9.57E-09 | Upstream | g10491 | 349 | CG11660 | serine/threonine-protein kinase RIO1 |
| 4 | 20445947 | 8.85E-09 | Downstream | g10649 | 17732 | | |
| 4 | 20445947 | 8.85E-09 | Upstream | g10648 | 5814 | | uncharacterized protein |
| 4 | 20445956 | 6.66E-10 | Downstream | g10649 | 17723 | | |
| 4 | 20445956 | 6.66E-10 | Upstream | g10648 | 5823 | | uncharacterized protein |

Table S3. GO categories showing enrichment in the GWAS hits.

| **Enrichment** | **FDR** | **nGenes Pathway** | **Genes Fold** | **Enrichment Pathways** | **GO ID** |
|---|---|---|---|---|---|
| 1.9E-03 | 3 | 106 | 49.4 | Positive reg. of nervous system development | GO:0051962 |
| 9.6E-04 | 4 | 215 | 32.5 | Reg. of nervous system development | GO:0051960 |
| 9.6E-04 | 4 | 242 | 28.9 | Positive reg. of developmental proc. | GO:0051094 |
| 5.4E-03 | 3 | 197 | 26.6 | Positive reg. of multicellular organismal proc. | GO:0051240 |

| | | | | | |
|---|---|---|---|---|---|
| 1.2E-03 | 4 | 281 | 24.9 | Reg. of multicellular organismal development | GO:2000026 |
| 7.6E-03 | 3 | 234 | 22.4 | Reg. of cell population proliferation | GO:0042127 |
| 3.3E-03 | 4 | 439 | 15.9 | Response to abiotic stimulus | GO:0009628 |
| 1.4E-03 | 5 | 691 | 12.6 | Reg. of developmental proc. | GO:0050793 |
| 5.9E-03 | 4 | 572 | 12.2 | Reg. of multicellular organismal proc. | GO:0051239 |
| 1.1E-03 | 6 | 1176 | 8.9 | Nervous system development | GO:0007399 |
| 2.0E-03 | 6 | 1443 | 7.3 | Cell development | GO:0048468 |
| 9.6E-04 | 7 | 1739 | 7 | Positive reg. of cellular proc. | GO:0048522 |
| 2.7E-03 | 6 | 1539 | 6.8 | System development | GO:0048731 |
| 9.6E-04 | 7 | 1892 | 6.5 | Positive reg. of biological proc. | GO:0048518 |
| 4.2E-03 | 6 | 1782 | 5.9 | Cell differentiation | GO:0030154 |
| 4.2E-03 | 6 | 1790 | 5.9 | Cellular developmental proc. | GO:0048869 |
| 4.2E-03 | 7 | 2831 | 4.3 | Anatomical structure development | GO:0048856 |
| 5.3E-03 | 7 | 3005 | 4.1 | Developmental proc. | GO:0032502 |
| 3.3E-03 | 8 | 4133 | 3.4 | Reg. of cellular proc. | GO:0050794 |

**SUPPLEMENTARY FIGURES**

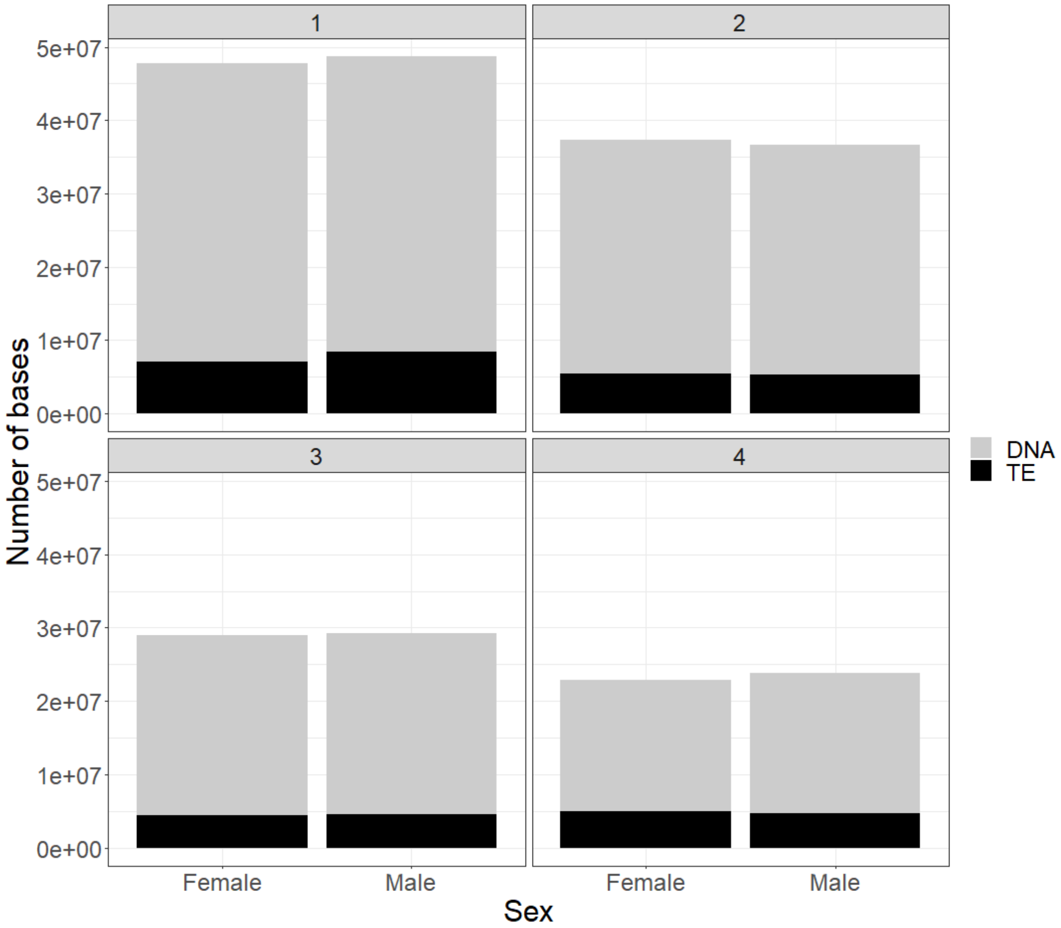


Figure S1. Overall number of bases classified as transposable elements (TE) and not classified as TE (DNA in figure legend). Facets represent the assembled chromosomes. Cumulative stacked bars represent total nucleotide bases per chromosome by sex.

Whole genome

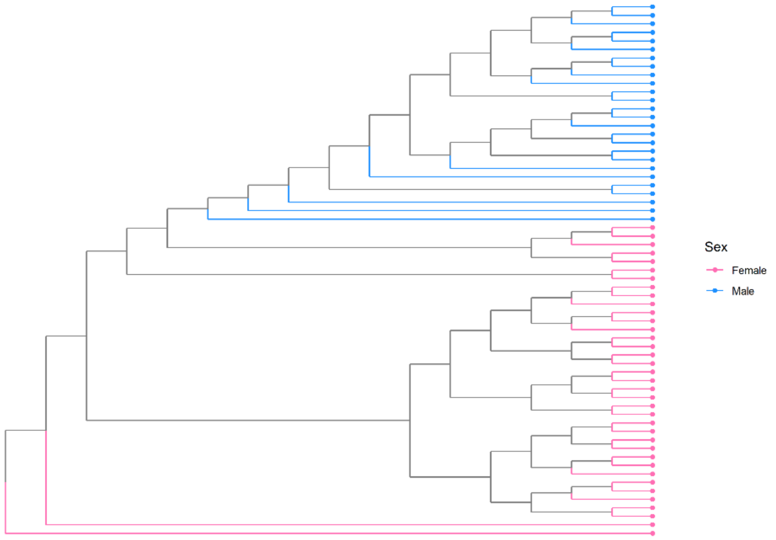


Chr1

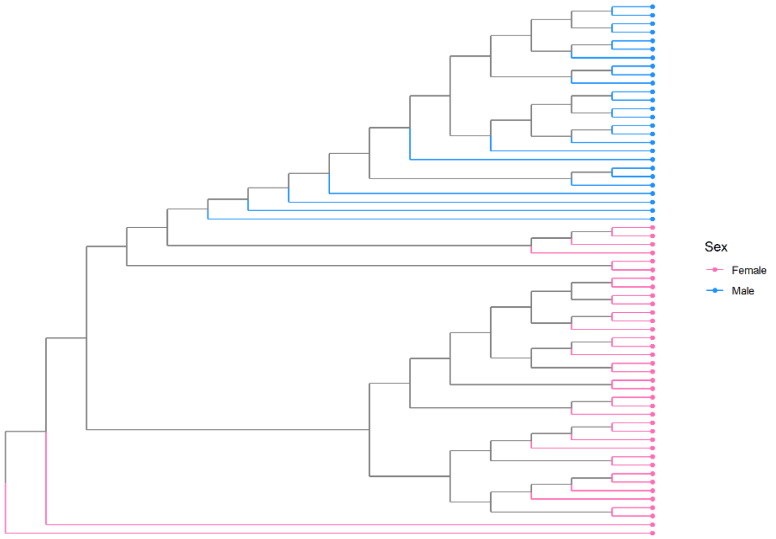


Autosomes

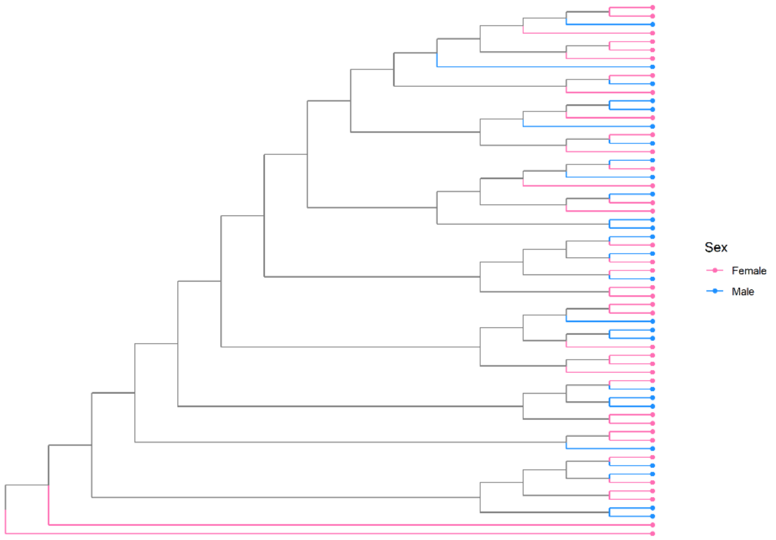

Figure S2. Phylogenetic trees of inter-relatedness between individuals across the whole genome, chromosome 1, and autosomes.

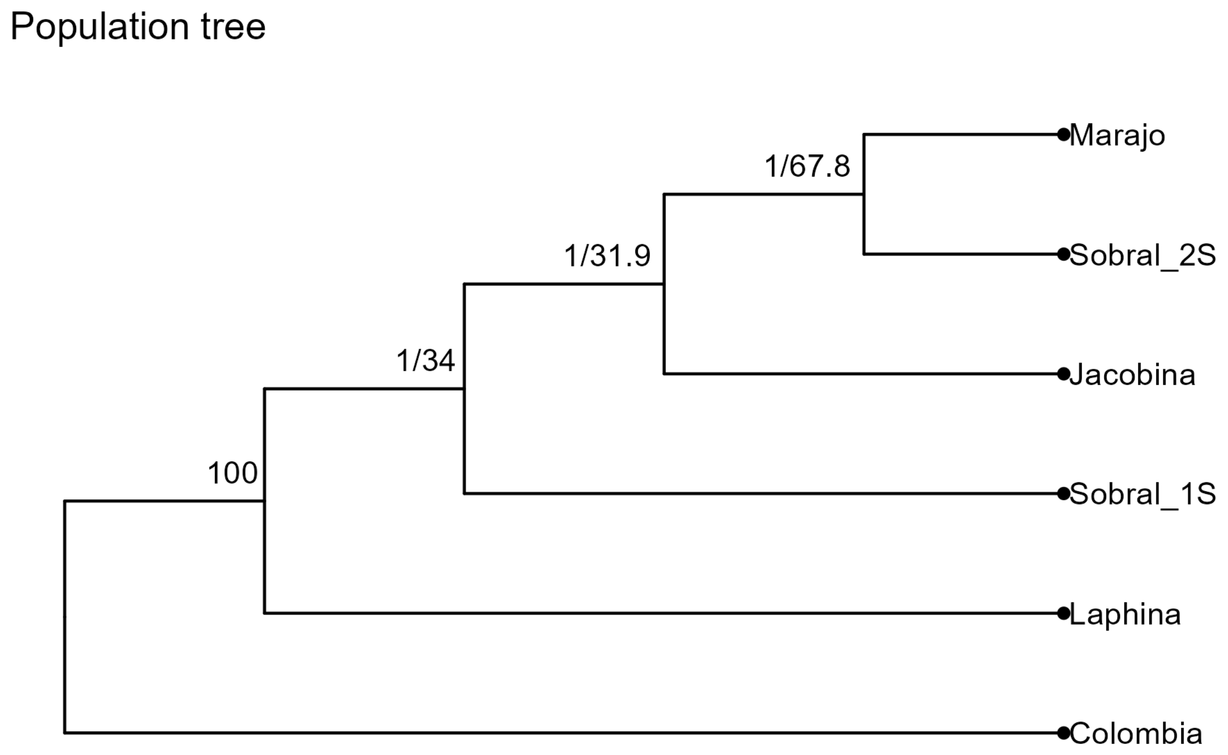


Figure S3. Consensus population tree from ASTRAL. The second values at the branches indicate gene concordance values from IQTREE2.

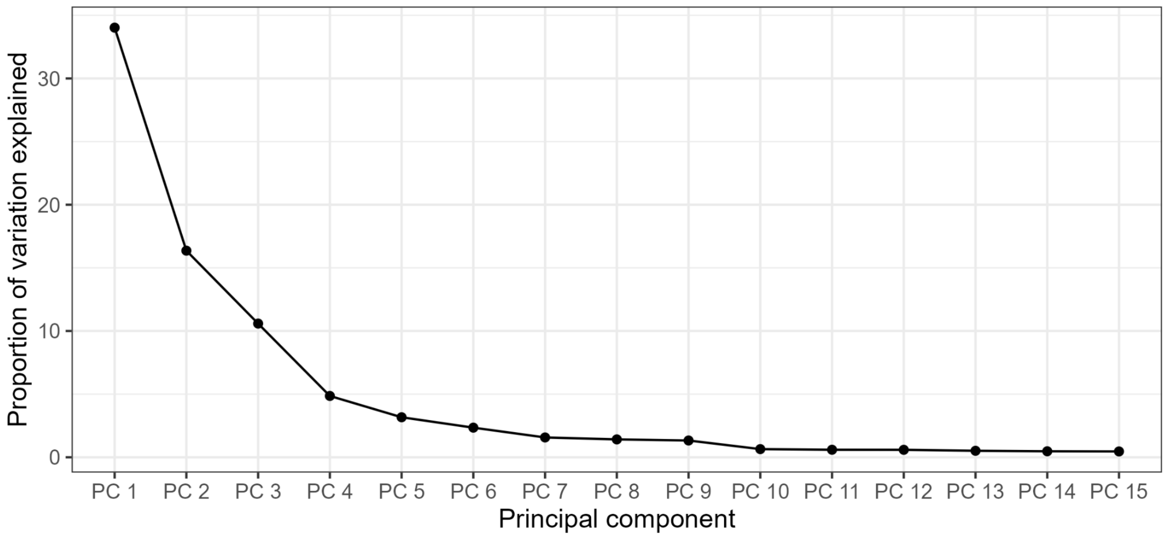


Figure S4. Scree plot of genomic variation in both Colombian and Brazilian sandflies explained. The proportion of variation explained is less than 3 after principal component 5.

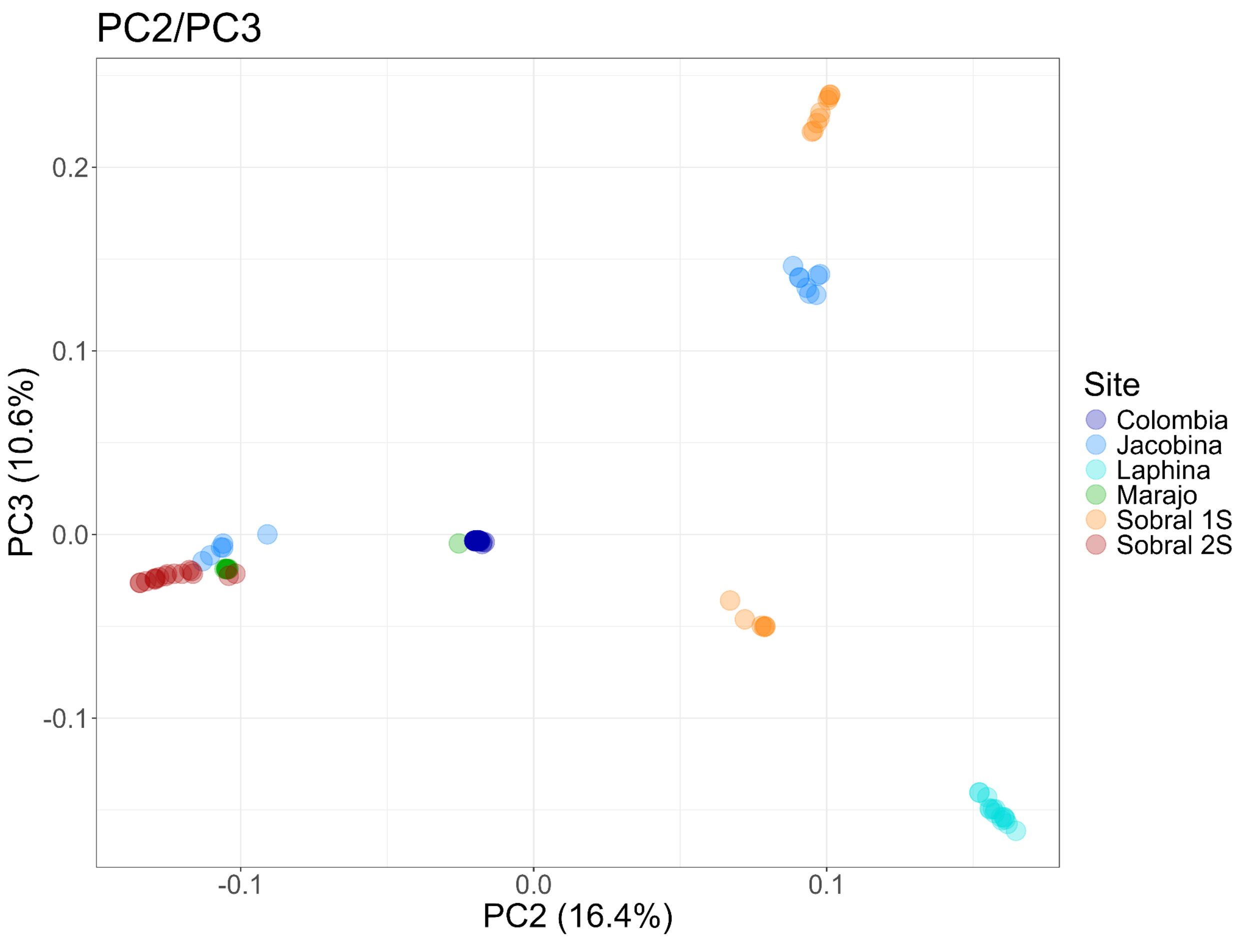


Figure S5. Principal components 2 and 3 from whole genome resequence data.

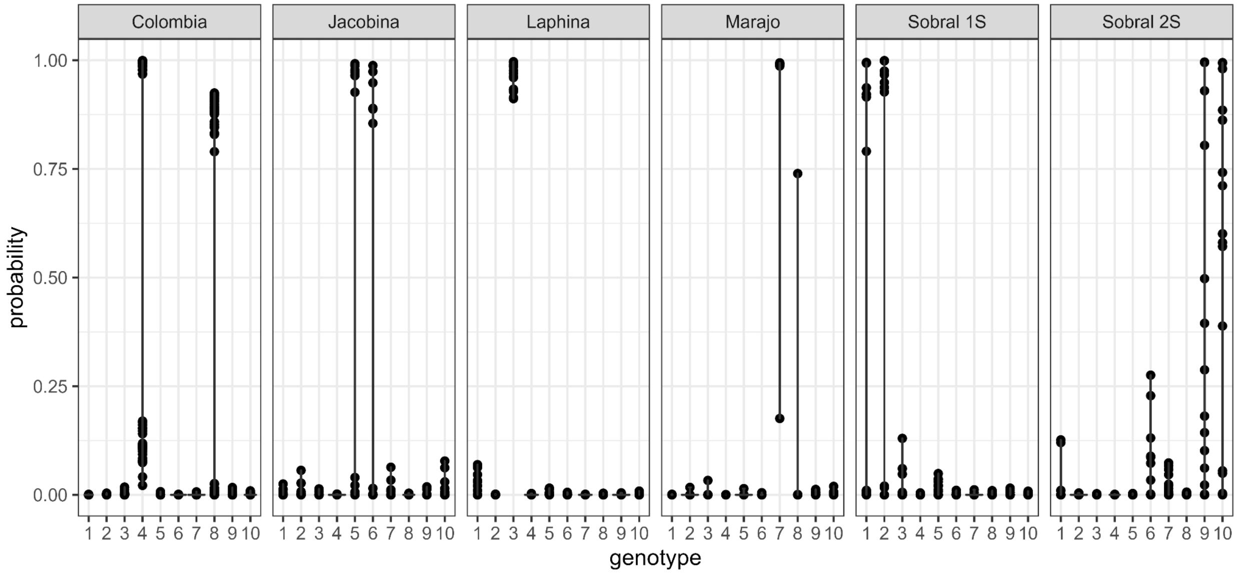


Figure S6. Admixture proportions by genotype for each individual sandfly.

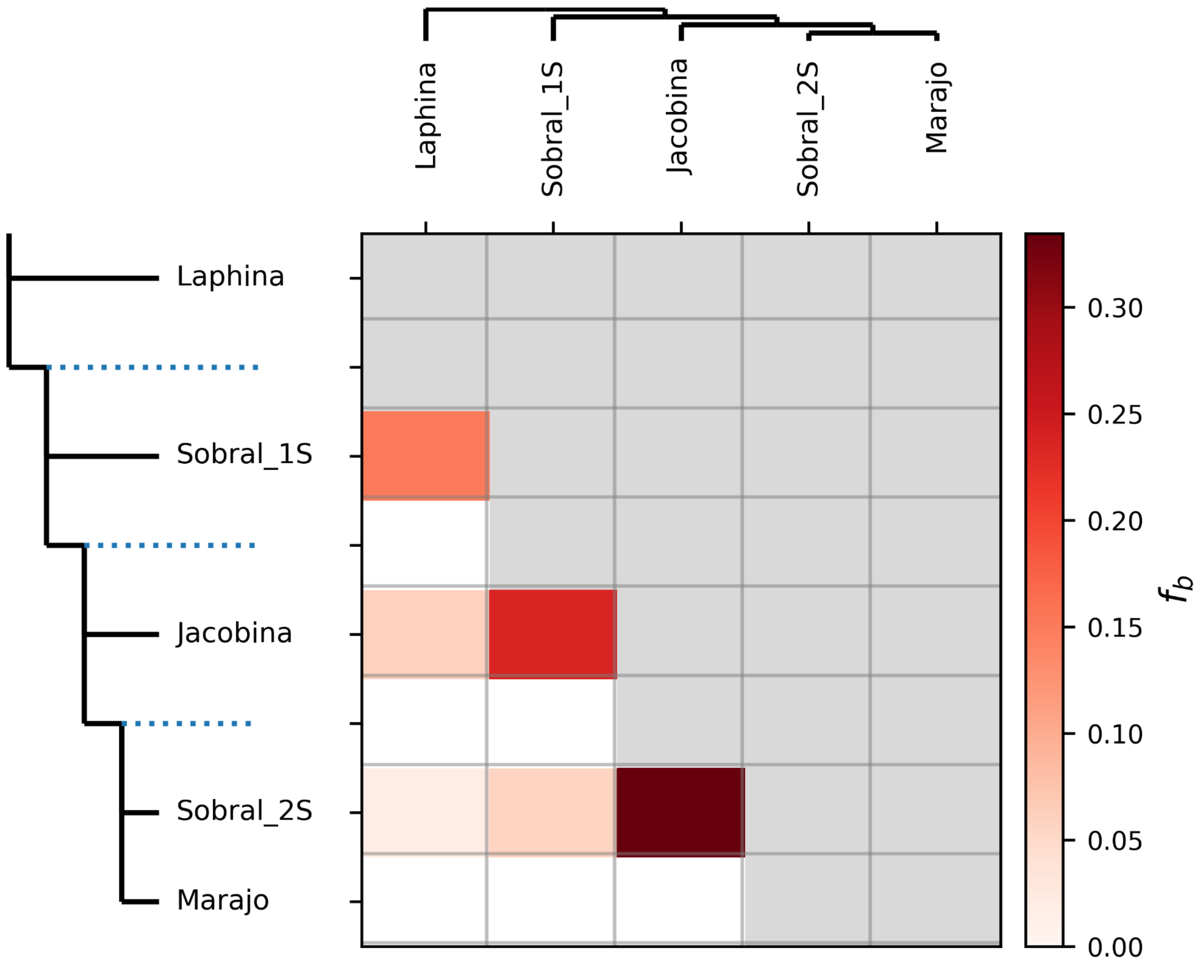


Figure S7. Fbranch statistics and gene flow inferred by Dsuite.

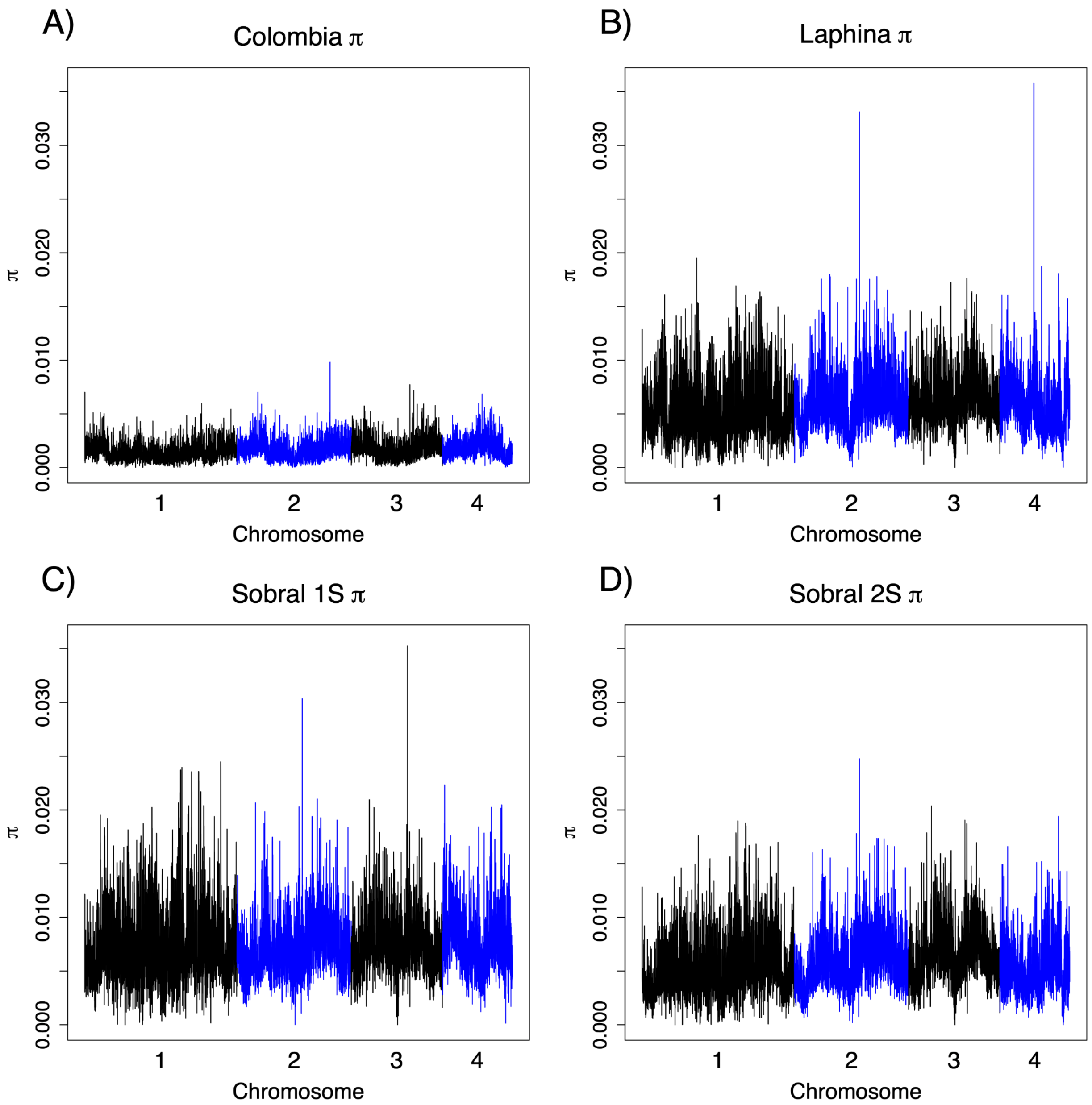


Figure S8. π in 10kb windows. **A.** π along the genome in Colombia. **B.** π along the genome in Laphina. **C.** π along the genome in Sobral 1S. **D.** π along the genome in Sobral 2S.

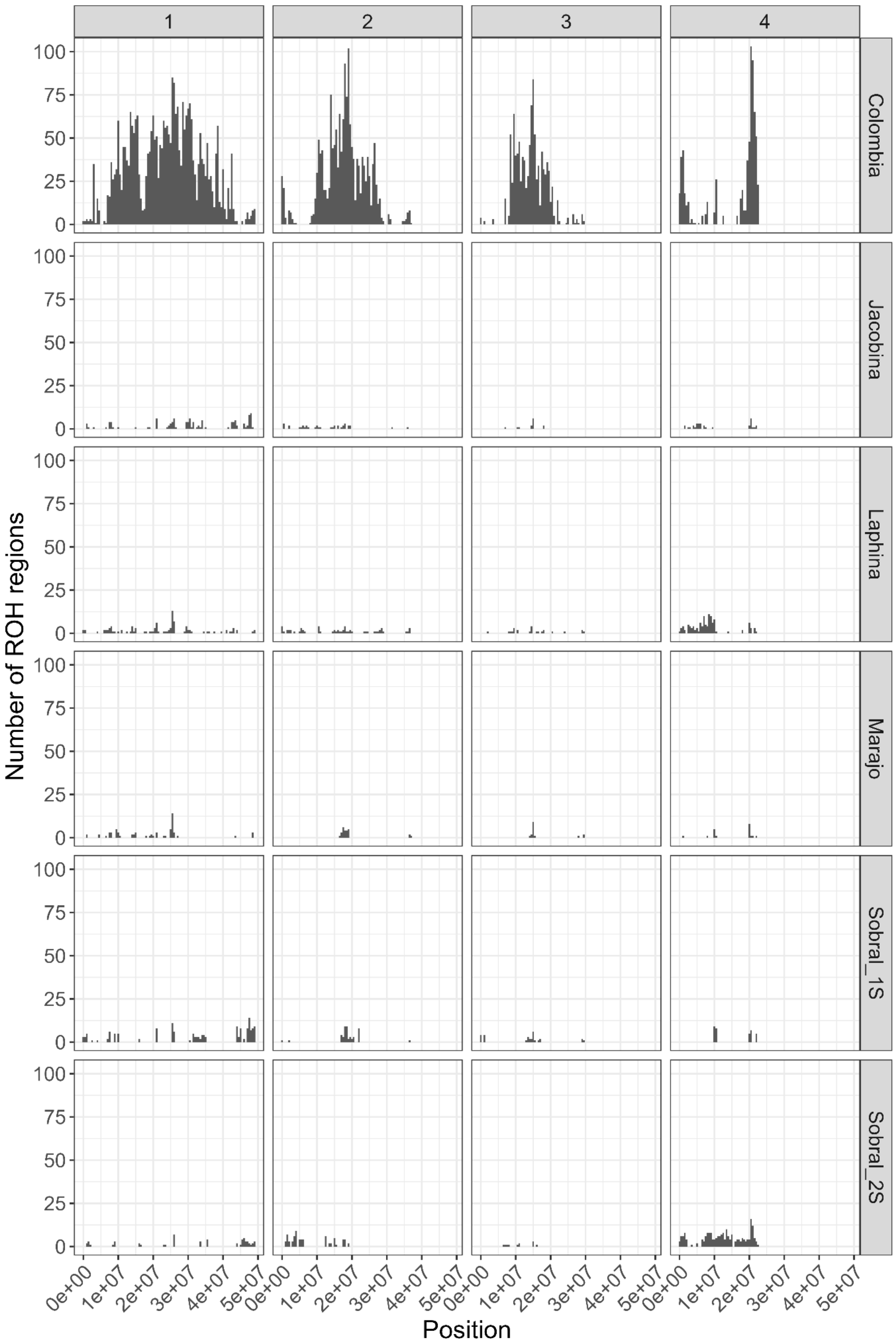

1
2
3
4
Colombia
Jacobina
Laphina
Marajo
Sobral_1S
Sobral_2S
Number of ROH regions
100
75
50
25
0
0e+00
1e+07
2e+07
3e+07
4e+07
5e+07
Position

Figure S9. Runs of Homozygosity in genomic regions from each population.

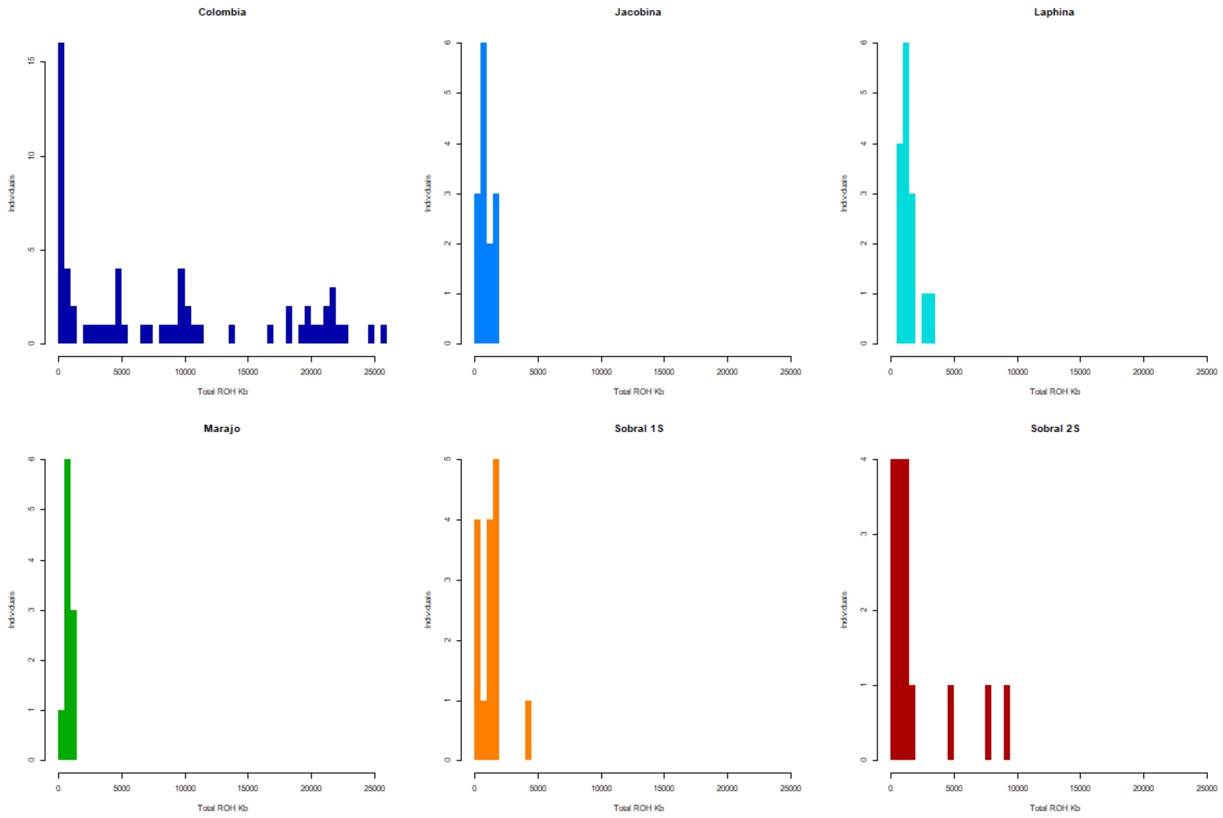


Figure S10. Histograms of ROH distributions by individual, clearly showing that the data from Colombia is not driven by a single individual.

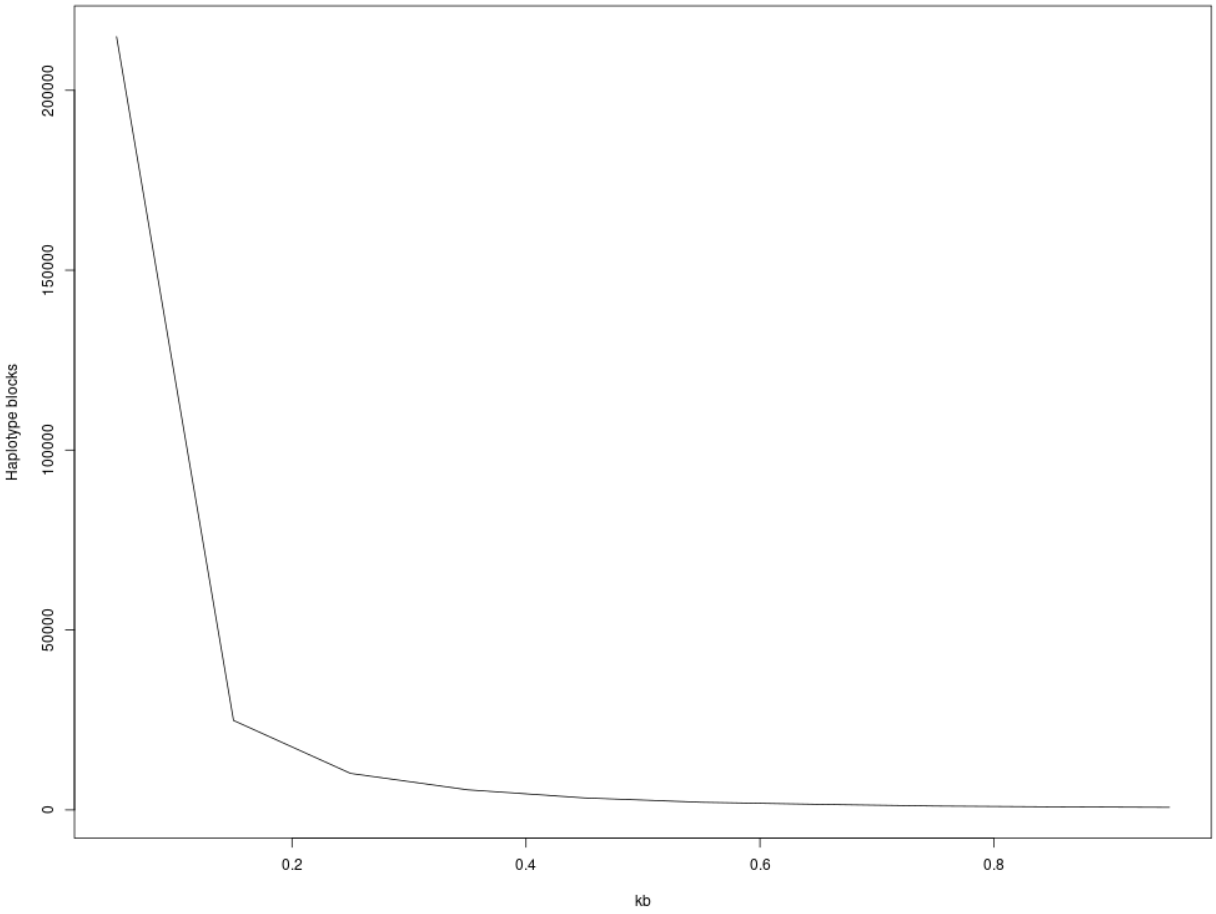


Figure S11. Linkage disequilibrium decays rapidly in *Lutzomyia longipalpis* with most haplotype blocks < 200bp.

DNA Extraction Protocol

Materials Needed:

- Zymo Genomic DNA Clean & Concentrator Kit
- Homogenization buffer (HB)
- 30mM Tris HCL (pH 8.0)
- 10mM EDTA
- 0:1M NaCl
- 0.5% Triton X-100
- SDS 10%
- RNase A
- Proteinase K

Note: This protocol is adapted from Dr. Jeremy Wang's high molecular weight extraction protocol for nanopore sequencing. Be sure that all solutions from the Zymo kit have the appropriate volume of 100% ethanol added.

1. Add 500 µL of HB to a 1:5mL tube with 20-40 frozen flies and then homogenize the flies using a clean pestle until you can no longer recognize any fly parts.
2. Add 125 µL of SDS and 10 µL of Proteinase K then vortex for 15 seconds.
3. Incubate at 55C for 1 hour (inverting every 10 minutes) and after 30 minutes add 10 µL of RNase A.
4. 4. Add twice the volume of ChIP DNA Binding Buffer to each volume of DNA sample and mix thoroughly.
5. Pipette the mixture into a Zymo-Spin Column in a collection tube, centrifuge at 16,000 x g for 30 seconds, and discard the flow-through.
6. Add 200 µL of DNA Wash Buffer to the Zymo-Spin Column, centrifuge at 16,000 x g for 1 minute, and discard the flow-through.
7. Repeat step 6.
8. Transfer the Zymo-Spin Column to a 1:5mL tube and to elute the DNA add 32 µL of DNA Elution Buffer directly onto the column matrix and centrifuge at 16,000 x g for 30 seconds.
9. Store the eluted DNA in the cold room until quantification.
